# Fusions of catalytically inactive RusA to FokI nuclease coupled with PNA enable programable site-specific double-stranded DNA breaks

**DOI:** 10.1101/2024.12.03.626739

**Authors:** Ahmed Saleh, Gundra Sivakrishna Rao, Qiaochu Wang, Magdy Mahfouz

**Author notes:** **Correspondence: Magdy M. Mahfouz**.

## Abstract

Programmable site-specific nucleases have revolutionized the genome editing. However, these systems still face challenges such as guide dependency, delivery issues, and off-target effects. Harnessing the natural functions of structure-guided nucleases offer promising alternatives for generating site-specific double-strand DNA breaks. Yet, structure-guided nucleases require precise reaction conditions and validation for *in-vivo* applicability. To address these limitations, we developed the <u>P</u>NA-<u>C</u>oupled <u>F</u>okI-(d)<u>R</u>us<u>A</u> (<u>PC-FIRA</u>) system. PC-FIRA combines the sequence-specific binding ability of peptide nucleic acids (PNAs) with the catalytic efficiency of FokI nuclease fused to a structurally-guided inactive RusA resolvase (FokI-(d)RusA). This system allows for precise double-strand DNA breaks without the constraints of existing site-specific nuclease and structure-guided nucleases. Through *in vitro* optimizations, we achieved high target specificity and cleavage efficiency. This included adjusting incubation temperature, buffer composition, ion concentration, and cleavage timing. Diverse DNA structures, such as Holliday Junctions, linear, and circular DNA, were tested demonstrating the potential activity on different target forms. Further investigation has revealed the PC-FIRA system capacity for facilitating the precise deletion of large DNA fragments. This can be useful in cloning, large-fragment DNA assembly, and genome engineering, with promising applications in biotechnology, medicine, agriculture, and synthetic biology.

## INTRODUCTION

Harnessing and redirecting natural molecular mechanisms have enabled understanding and manipulating the genetic code of life. Site-specific nuclease (SSNs) such as zinc-finger nucleases (ZFNs), transcription activator-like effector nucleases (TALENs), and clustered regularly interspaced palindromic repeats (CRISPR)-associated proteins (Cas) facilitated a rapid expansion beyond basic biology research to applications in diagnostics, genome engineering, biosynthesis, and therapeutics *^1-4^*. Despite its broad potential, CRISPR-Cas technology present limitations such as requiring the presence of a 2–6 bp protospacer adjacent motif (PAM), hindering its use in PAM-independent applications *^5^*. Additionally, limitations such as large size, unstable and charged single guide RNA (sgRNA), and cell membrane barriers constrain its delivery *^6, 7^*. Exploring tools beyond CRISPR-Cas can offer novel approaches, avoiding any persisting limitations. For instance, the recently developed PNA-assisted pAgo editing (PNP editing) system have exploited the natural function of prokaryotic argonautes (pAgos) to induce programmable targeted double-strand DNA breaks (DSBs) *^8, 9^*. PNP editing has demonstrated significant potential in target DNA cleavage in vitro. However, PNP editors are still to be investigated thoroughly for intended target application in vivo.

Peptide nucleic acids (PNAs) are synthetic nucleic acid analogs *^10^*. Unlike DNA, PNAs have a pseudopeptide backbone, which allows them to form sequence-specific and stable bonds with both single-strand DNA (ssDNA), RNA and double-strand DNA (dsDNA) through a process called invasion. Additionally, PNAs are highly resistant at cellular conditions against any nucleases or proteases due to their artificial electrostatically neutral pseudo-peptide backbone. These unique characteristics make PNAs a highly valuable tool for DNA targeting applications *in vitro* and *in vivo ^11^*. For instance, the use of PNAs in DNA editing systems has shown promise, for example, γtcPNA-mediated gene editing was used to correct anemia in β-thalassemic mice i*n vivo ^12^*. Due to their strand invasion behavior, they have been implemented as probes in a SARS-CoV-2 detection system, and as DNA-openers in the PNP editing system *^9, 13^*.

Recently, we established two different DNA manipulation systems called PNA-assisted resolvase mediated (PNR) editing and PNA-guided T7EI (PG-T7EI) editing, expanding on the exploitation of PNAs and resolvases. Resolvases are highly specialized, in nature mainly recognizing and resolving Holliday Junctions (HJs) that form during DNA repair and genetic recombination *^14, 15^*. By redirecting the functions of PNAs to invade target DNA and to simulate HJ analogs that are recognizable by structurally-guided resolvases such as T7 Endonuclease I (T7EI) and *Arabidopsis thaliana* MOCI (AtMOCI), we developed efficient, precise and target specific dsDNA cleavage tools *^16-19^*. These findings suggest that structure guided nucleases (SGNs) could be used to create customizable, efficient DNA manipulation tools that address challenges posed by current SSNs. For instance, the developed PG-T7EI editing tool was able to withstand a wide range of reaction conditions and facilitate cloning of large DNA fragments *^18^*. Nevertheless, SGNs have revealed their limitations with some resolvases lacking structure specificity indicating possible off-target activity *^18, 19^*. Despite these challenges, SGNs remain a promising option, offering a massive library of novel proteins to be explored for their potential in DNA editing applications *^20^*.

RusA resolvase, a 14 kDa homodimer endonuclease encoded by a cryptic prophage gene, is conserved in many species, including *Escherichia coli*, *Aquifex aeolicus*, *Mycoplasma pneumoniae*, and various other endosymbiotic bacteria *^21^*. The natural function of RusA provides an alternative pathway for HJ resolution when RuvC or the entire RuvABC complex is nonfunctional, highlighting its role in cell survival *^22, 23^*. RusA resolves naturally occurring and synthetic HJs by binding and cleaving HJs in a structure-specific and sequence-specific manner, respectively *^21^*. RusA first recognizes and binds to X-form four-way junctions, then induces a Mg^2+^ dependent dual-strand cleavage at the (5′-▾CC-3′) dinucleotide located symmetrically at the junction core. Previous studies used site-directed mutagenesis to identify specific amino acid conversions crucial for catalytic activity, including R69A, K76A, and D70N *^24^*. Of these, D70N was found to be the most critical, as the original aspartic acid plays a key role in coordinating the metal ion interactions necessary for catalysis. RusA has been used as a eukaryotic genome probe with high selectivity for HJs, operating independently of other replication machinery, highlighting its functionality *in vitro* and *in vivo ^24-26^*.

FokI nuclease is a key genome engineering tool due to its compact size, structural properties and cleavage activity *^27^*. It has been used to enhance the editing efficiency of SSNs and also for enhancing specificity through dimer formation. For instance, in CRISPR-Cas technology, FokI was fused to catalytically dead Cas9 (dCas9), achieving >130 fold target specificity compared to wild-type Cas9 *^28^*. Additionally, PNA-assisted FokI-(d)pAgo editors (PNFP editors), an extension of PNP editing, involved fusing FokI to catalytically dead pAgos, adding an extra layer of specificity to PNP editing *^9, 29^*.

We identified RusA resolvase as an ideal candidate for developing a more precise and efficient DNA editing system. Inspired by PNR-editing, we hypothesized that by exploiting the structural ability of dead RusA ((d)RusA) to bind selectively to four-way HJs simulated through PNAs sequence specific dsDNA invasion, and through the catalytic activity of FokI nuclease fused to (d)RusA, we could induce programmable accurate and precise DSBs. We termed this novel system, PNA-Coupled FokI-(d)RusA (PC-FIRA). To validate our theory, we tested Wildtype RusA ((wt)RusA), (d)RusA, and FokI-dead RusA (FokI-(d)RusA) proteins on different target forms and confirmed that only FokI-(d)RusA fusion is able to bind to the PNA-simulated HJ-analogs and induce cleavage. Furthermore, we tested the cleavage reaction under different conditions in vitro to identify optimal reaction conditions in which we can generate accurate and precise dsDNA DSBs. In addition, we also investigated the potential of the PC-FIRA system in manipulating large fragments release through multiplexing cleavage.

## METHODS

### RusA cloning and purification

Protein structure predictions for (wt)RusA, (d)RusA, FokI- (d)RusA and (d)RusA-FokI were performed to infer protein conformations and stability considering amino acid modifications and fusions. FokI was placed at both N and C terminal of RusA through a 33 amino acid linker. *In silico* prediction was performed using Alphafold 3.0 (https://alphafoldserver.com) *^30^* for (wt)RusA, (d)RusA, FokI-(d)RusA, and (d)RusA-FokI. All the protein sequences are listed in Table S1.

All the gene sequences were codon-optimized for *E. coli*, then ordered as G-Blocks from Integrated DNA technologies, Inc. G-blocks were further PCR-amplified using a gene specific forward and reverse primers containing BamHI and XhoI restriction enzymes, respectively (Table S6). The PCR products were purified from a 1% agarose gel, cloned into pET28a bacterial expression plasmid under 6xHis-tag (Files S1). Clones were confirmed through Sanger sequencing, and introduced into chemically competent BL21-PLYSS(DE3) *E. coli* strain.

For protein purification, a single colony was inoculated into 100 ml Luria-Bertani (LB) broth with 50 mg/ml kanamycin and incubated for 16 h at 37°C and 170 rpm. Then, 20 ml from the overnight grown culture was inoculated into one-liter Terrific Broth (TB) (IBI SCIENTIFIC, 23H3013) liquid media (51g TB and 8ml 50% glycerol, 50 mg/ml kanamycin). Likewise, we prepared four one-liter flasks. The one-liter cultures were incubated at 37°C until OD reached 0.5 at OD600. Cell growth was slowed by instantly placing the cultures at 4°C for 30 mins with shaking to ensure cell resuspension. Protein expression was induced by adding 0.25 mM Isopropyl β-d-1-thiogalactopyranoside (IPTG), and the cultures were incubated for 16 h at 18 °C and 150 rpm. Induced cultures were collected independently into four one-liter centrifuge tubes then centrifuged for 20 mins at 5000 rpm using a Thermo Scientific Sorvall LYNX 4000 centrifuge.

The cell pellet was then resuspended in lysis buffer (4 ml buffer for 1g of cell pellet) (Table S2) using a magnetic bead for 30-40 mins at 4°C. Then the lysate was sonicated at 4°C using a Qsonica Q700 sonicator with 5-sec ON and 10-sec OFF intervals at an amplitude of 40. The cell lysate was then centrifuged at 4°C for 20 mins at 16000 rpm and filtered using a 0.45-μm flow bottle top filter (Thermo Scientific Nalgene, 291-4545).

To capture the desired protein, the lysate was passed through an affinity chromatography column (HisTrap HP, 5 ml, GE Healthcare) in ÄKTA pure 25 M1 protein purification system (Max flow rate 25 mL/min, pressure range 0–20 MPa, multi-wavelength detection 190–700 nm), then eluted the protein using a high imidazole buffer (50 mM Tris–HCl at pH 7.5, 500 mM NaCl, 300 mM Imidazole, 1 mM TCEP, and 5% glycerol). All the collected fractions were run on an SDS gel and the correct protein fractions were then dialyzed overnight (∼16 hrs) in dialysis buffer (25 mM Tris–HCl at pH 7.5, 100 mM NaCl, 1 mM TCEP, and 5% glycerol). A second protein purification step was performed using a cation-exchange column (HiTrap SP HP, 5 ml, GE Healthcare) with a high salt elution buffer (50 mM Tris–HCl at pH 7.5, 2 M NaCl, and 1 mM TCEP). The collected fractions were run on an SDS gel for purity check and the selected fractions were processed for high salt removal. Buffer exchange was done using Amicon Ultra-15 centrifugal filter units (10 kDa NMWL, Millipore) with a storage buffer (25 mM Tris–HCl, 100 mM NaCl, 1 mM TCEP, and 10% glycerol). Based on protein concentration measured with a NanoDrop Eight spectrophotometer, the collected fractions were concentrated to 1.3, 0.6 and 0.05 mg per ml for (wt)RusA, (d)RusA, and FokI-(d)RusA, respectively. Finally, aliquots were collected, snap-frozen, and stored at -80°C.

### Design and synthesis of γPNAs

The PNA used in this work were designed and tested in previous studies showing enhanced binding affinity, specificity, and stability by modifying all γ-positions with the amino acid alanine and flanking three lysine residues at each end (Table S4) *^18,19^*. General PNA synthesis guidelines (https://www.pnabio.com/support/PNA_Tool.htm) were followed, and the γPNA were synthesized by HLB PANAGENE Co., Ltd. Republic of Korea.

### Design and cloning of PNA-binding pUC19 targets

Fragments with target sequences for γPNA1 were ordered from Integrated DNA Technologies, Inc. as top and bottom oligos, which were phosphorylated, annealed, producing overhangs that allow cloning by ligation into NdeI-digested pUC19 (Table S5). The cloned sequences were confirmed by both restriction analysis and Sanger sequencing using forward and reverse primers, and positive clones were later propagated for future use (Table S6). The cloned plasmid was named as pUC19-γPNA1-151. The γPNA1 binding sequences in the pUC19-γPNA1-151 are 20 bp length each and separated by a 13 bp spacer (Files S2a).

### Design and synthesis of Holliday Junctions

Four separate oligos from Integrated DNA Technologies were ordered for producing the HJ structures. The four oligos, 1ul of each oligo from 1 µM stock, were annealed in equimolar concentrations in a 10ul reaction. The produced HJs were used to examine (wt)RusA and (d)RusA activity. The activity assays were adopted from previous work *^31^*. Additional complementary flanking sequences of approximately 45 bases were added to each strand of the adopted core homology sequence, forming a 200 bp junction that can be observed on the gel (Table S3). The junction involves **D**ouble **C**leavage of strands **1** and **3** (DC-1&3) at the core homology sequence containing two dinucleotides (5′-▾CC-3′) positioned diagonally. DC-1&3 contains target sites for PvuII and SphI on strands 1&4 and 2&3, respectively. These two enzymes were used as positive restriction control, each cutting away 21 and 17 bp fragments respectively and forming a truncated HJ with a size around 160 bp.

### γPNA invasion and RusA-based restriction

Invasions of circular and XbaI-linearized pUC19-γPNA1-151 targets were performed with 150 nM γPNA and 50 ng dsDNA (3960bp) in 1X MOPS buffer (20 mM MOPS, pH 7.0, 5 mM CH3COONa, and 1 mM EDTA), making up a total reaction volume of 10 μL. The invasions were incubated at 37°C for 2 hrs and 6 hrs, corresponding to circular and linear target invasions, respectively. Cleavage reactions with (wt)RusA, (d)RusA, and FokI-(d)RusA were standardized and assembled as mentioned in the Table 1.

**Table 1.**
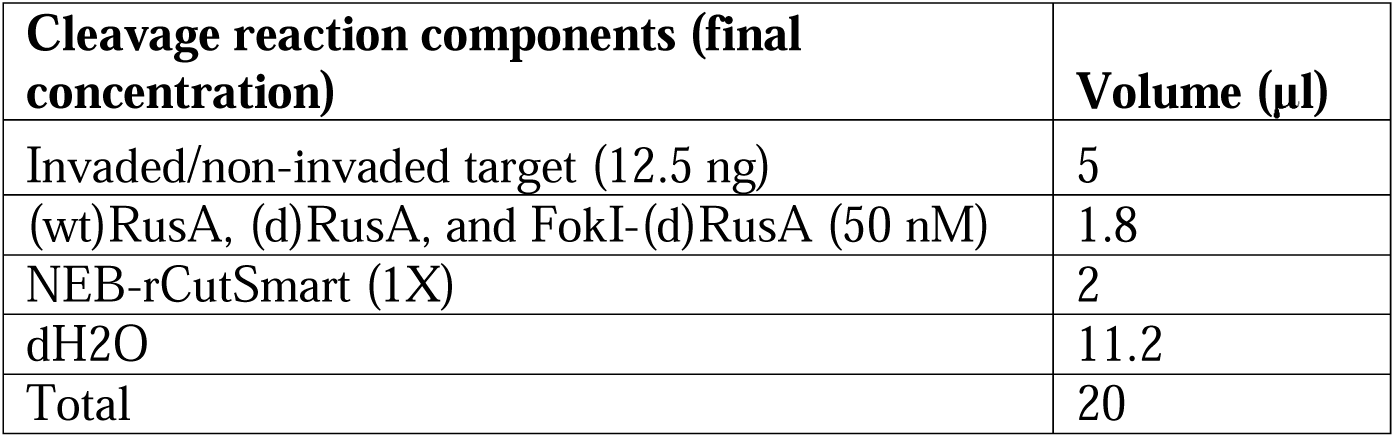
γPNA invasion and RusA based restriction reaction components and concentration.

80 mM NaCl was added to all FokI-(d)RusA reactions, and 2mM MgCl2 was added to (wt)RusA reactions to test its effect on (wt)RusA activity. XbaI or XmnI restriction enzymes were used to linearize the pUC19-γPNA1-151 target or added to the (wt)RusA, (d)RusA, and FokI-(d)RusA cleavage reaction to ensure the band release. dH_2_O volume was adjusted depending on the addition of restriction enzymes, salts, and multiple PNA invasions. Samples were incubated for one hour, then 5 μL of 6X Purple Loading Dye (NEB, B7024S) were added and run on a GelRed-stained 1% (w/v) agarose gel for 2 h at 150V. Finally, the gel was visualized using an iBright 1500 imaging system (Thermo Scientific, A44114). All restriction enzyme-treated and untreated samples were subjected to 37°C for 15 minutes.

Almost all experiments followed the aforementioned reaction conditions; however, minor modifications to volumes and reagents were applied based on experiment requirements, such as performing protein concentration titration, using different salts, buffers, and temperatures, and conducting cleavage over different time points.

### FokI-(d)RusA mediated multiplex cleavage of linear targets

To explore the feasibility of utilizing FokI-(d)RusA for larger fragment excision by multiplex cleavage of γPNA-invaded targets, we designed a target plasmid cloned with two γPNA1 target sites separated by a 1291 bp spacer (Files S2b). The plasmid pUC19-γPNA1&1-245 was first linearized with XmnI, then invaded by γPNA1 and treated with 50 nM FokI-(d)RusA and 80 mM NaCl. The reaction was incubated at 30°C for 60 minutes. BsrGI and BamHI were used as restriction control on non-invaded XmnI-linearized pUC19-γPNA1&1-245 target, with both enzymes corresponding to the two γPNA1 target sites. The expected cleavage products predicted for BamHI is 1917 bp, and 1881 bp fragments; for BsrGI is 3195 bp and 603 bp; and for BamHI and BsrGI together is 1881 bp, 1314 bp, and 603 bp.

## RESULTS AND DISCUSSION

### PC-FIRA system overview

Developing a genome engineering tool that balances between specificity, efficiency, deliverability, minimal off-target effects, and versatility remains a major challenge. Even transformative tools like CRISPR-Cas face limitations constraining their utility in clinical and therapeutic applications. Here, we propose the PC-FIRA system as an efficient and streamlined DNA manipulation tool. This system is designed to overcome some of the challenges associated with existing technologies, providing a complementary or alternative approach to genome modification.

The PC-FIRA system leverages the structural specificity of RusA for the binding of HJ kind of DNA structure generated by PNAs, while bypassing its inherent sequence specificity constraints for DNA cleavage. The catalytic domain of FokI, known for its efficient nuclease activity, facilitates precise and controlled DSBs (Figure 1a). *In vitro* assays have demonstrated the capacity of PC-FIRA to induce targeted DNA cleavage; however, this activity is dependent on optimized reaction conditions that maximize both specificity and efficiency. Through systematic optimization of parameters such as temperature, ion concentration, and incubation times, we identified conditions that promote optimal catalytic activity of the FokI-(d)RusA fusion protein, thereby enhancing its potential as a versatile tool for genome editing applications.

**Figure 1.**
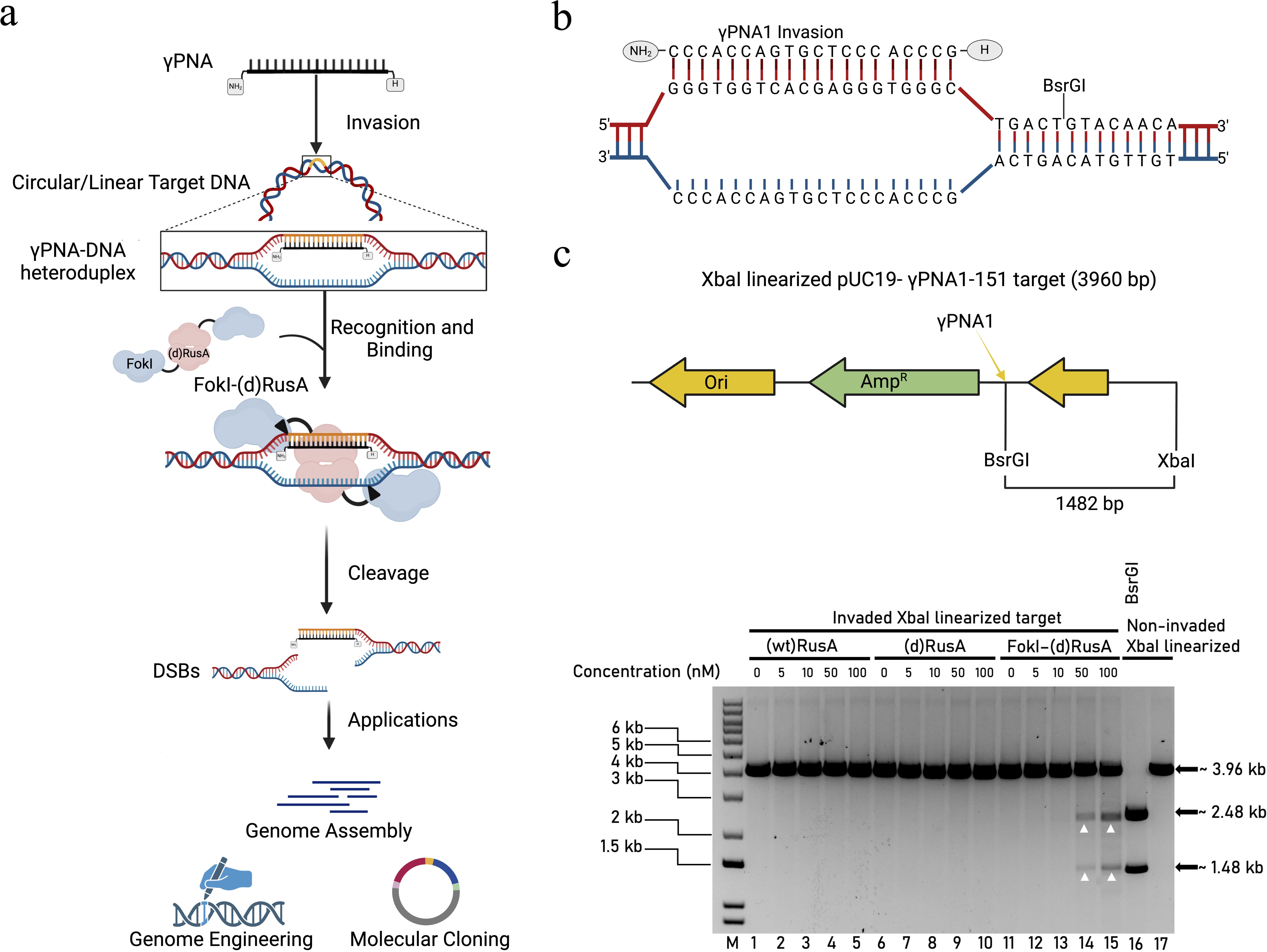
Scheme of PC-FIRA system and cleavage activity of (wt)Rus, (d)RusA, and FokI-(d)RusA under optimal conditions on invaded linear target. (a) Sketch showing γPNA invasion, FokI-(d)RusA target recognition and dsDNA break. The potential applications of PC-FIRA after dsDNA break are shown. (b) Sequence information of γPNA1 invaded at target region in pUC19-γPNA1-151 plasmid. (c) XbaI linearized pUC19-γPNA1-151 plasmid map showing γPNA1 invasion region and BsrGI restriction enzyme site. Gel image shows the cleavage activity of (wt)RusA (Lanes 1-5), (d)RusA (Lanes 6-10), and FokI-(d)RusA (Lanes 11-15) at increasing protein concentrations on γPNA1 invaded XbaI-linearized pUC19-γPNA1-151 target plasmid. FokI-(d)RusA cleavage products are indicated by arrowheads. BsrGI and XbaI restriction enzymes cleaved and XbaI-linearized non-invaded pUC19-γPNA1-151 was included in lane 16 and 17, respectively as positive size controls. Lane M shows the 1-kb plus DNA marker.

### Protein structure stability and initial activity assessment

Using AlphaFold to predict the structure of (wt)RusA, (d)RusA, and FokI-(d)RusA, we inferred preliminary insights on the stability and the integrity of FokI and (d)RusA fusions. Placing FokI at either the C or N - terminal of (d)RusA might influence activity and stability of the fusion due to domains interference. Furthermore, the length and composition of linkers fusing FokI to (d)RusA can have an impact on function. Short rigid linkers may restrict FokI movement, altering its catalytic efficiency, and long flexible linkers could permit FokI-independent activity, leading to undesired cleavage activity*^32^*. AlphaFold results revealed >90 score after predicted Local Distance Difference Test (pLDDT) for both (d)RusA and an N-terminal FokI fused (d)RusA. We assured that the target recognition and target binding ability of (d)RusA is not influenced by N-terminal fused FokI nuclease. In addition, any steric influence of (d)RusA on FokI catalytic activity is also avoided by using a 33-amino acid linker. AlphaFold results also suggested a well-folded and stable protein with low expected positional error (Figure S1).

The purified (wt)RusA, (d)RusA, and FokI-(d)RusA proteins were run on an SDS-PAGE gel with increasing concentrations, (wt)RusA (15, 30, and 50 µM); (d)RusA (6, 12, and 24 µM); and FokI-(d)RusA (0.4, 0.8, and 1.2 µM) to check for any minor impurities. Qualitative assessment of the gel confirmed that all proteins were pure enough to proceed for further characterizations (Figure S2).

We initially tested the activity of (wt)RusA and (d)RusA on a designed HJ named Double Cleavage Strand 1 and 3 (DC-1&3) which allows for proper binding of dimerized RusA then correctly positioning of the dinucleotide within the core homology sequence at the cleavage site of RusA for cleavage (Figure S3a). Almost complete cleavage was noticed with DC-1&3 treated with 50 nM (wt)RusA, producing two 100 bp fragments observed on a 3% agarose gel, confirming (wt)RusA activity (Figure S3b). Untreated and 50nM (d)RusA treated DC-1&3 showed slightly higher band size than 200 bp which is expected due to HJ slower migration. Positive control samples with restriction enzymes showed a higher shift in band size, confirming the excision of fragments from DC-1&3 arms forming a truncated HJ. In addition, it was also observed that in the absence of MgCl2, (wt)RusA was still able to induce cleavage on DC-1&3, which could be mediated by Mg^2+^ supplied with rCutsmart.

### PC-FIRA mediated precise target cleavage

By establishing optimized cleavage reaction conditions, we were able to generate controlled accurate DSBs using FokI-(d)RusA. Initially, the XbaI restriction enzyme linearized pUC19-γPNA1-151 target was invaded with γPNA1 molecule (Figure 1b). The γPNA1 invaded linear pUC19-γPNA1-151 target was treated with increasing concentrations (0nM, 5nM, 10nM, 50nM, and 100 nM) of (wt)RusA, (d)RusA, and FokI- (d)RusA. Protocol for optimal reaction conditions is presented in the methods. No binding or cleavage activity for (wt)RusA and (d)RusA were observed on γPNA1 invaded linear targets in optimal conditions. Whereas, the FokI-(d)RusA showed significant targeted cleavage activity at 50 nM and 100 nM with cleavage efficiencies of 16% and 32% calculated using ImageJ, respectively (Figure 1c). In the same optimal reaction conditions, (wt)RusA, (d)RusA, and FokI- (d)RusA did not show any target cleavage or specific target cleavage of γPNA1 invaded circular pUC19-γPNA1-151 target (Figure S4).

### PC-FIRA activity on different forms of the target plasmid

Recent PNFP editors work has indicated that FokI fusions do not require the invasion of a pair of PNAs for target sites in close proximity to mediate cleavage activity *^29^*. Furthermore, considering that RusA is present as a homodimer within a solution, we assumed that a single γPNA molecule invasion can simulate a four-way HJ structure and allow recognition, binding, and cleavage.

Non-invaded and γPNA1-invaded, circular and linear pUC19-γPNA1-151 targets were treated with (wt)RusA, (d)RusA, FokI-(d)RusA, and FokI, independently at increasing concentrations (0, 5, 10, 50, 100, 200 nM). (wt)RusA and (d)RusA treated targets showed no cleavage activity on both non-invaded and invaded circular and linear targets (Figures S5a and S5b; Figures S6a and S6b). γPNA1 invaded linear and circular pUC19-γPNA1-151 treated with FokI-(d)RusA showed a band of interest including non-specific cleavage. The non-specific cleavage occurred more on circular target (Figure S7a and S7b). Non-invaded circular and linear targets treated with FokI-(d)RusA showed non-specific cleavage and no release of band of interest (Figure S8a and S8b). FokI showed high non-specific cleavage and degradation of circular products at concentrations ≥ 50 nM, which is higher compared to FokI-(d)RusA with degradation starting at 200 nM.

These findings suggest that (wt)RusA cannot function effectively on PNA-invaded DNA targets that do not possess the sequence specificity at the preferred position at HJ kind of structures for mediating the cleavage. This suppressed inactivity adds a layer of specificity in cells possessing naturally occurring (wt)RusA. However, in the current system, we exploited only the structural specificity of (d)RusA and the catalytic activity of FokI contributing to specific binding and cleavage, respectively. This approach enhances the versatility of the system as a genome editing tool.

Notably, the non-specific cleavage observed by FokI-(d)RusA is significantly reduced on linearized invaded targets compared to circular invaded targets. This difference is likely due to the supercoiled nature of circular DNA. Circular DNA can form symmetrical structures that resemble HJs, and this structural similarity might be a contributing factor to non-specific cleavage *^33^*. Understanding the order of binding and dimerization of RusA and FokI within the PC-FIRA system could potentially provide insights on reducing non-specific activity. Linear genomic DNA is the predominant form in eukaryotic cells, the FokI-(d)RusA fusion protein could be particularly advantageous for genome editing applications that favor linear DNA targets, thereby potentially minimizing off-target effects. Furthermore, considering the previous success using (wt)RusA as a probe in yeast, it seems plausible to investigate the applicability of our PC-FIRA system for broader *in-vivo* applications, offering exciting possibilities for further research.

### Optimization of ion concentration

Previous work indicated that the use of NaCl at certain concentrations in cleavage reactions could lower non-specific cleavage activity *^29^*. This is likely due to the stabilization of specific DNA-protein interactions and the disruption of weaker non-specific interactions, as higher ionic strength mitigates electrostatic repulsions between the negatively charged DNA and the enzyme *^34^*.

Hence, we tested the cleavage activity of FokI-(d)RusA in the presence of NaCl aiming to determine the optimal ionic strength for minimizing non-specific cleavage while maintaining efficient enzyme activity. FokI-(d)RusA activity was assessed across varying NaCl concentrations (0, 10, 20, 30, 40, 60, 80, 100, 110, 130, 150 mM) on γPNA1 invaded XbaI-linearized pUC19-γPNA1-151 (Figure 2). Results demonstrated that non-specific cleavage with linear target was greatly reduced, and the clean released bands were observed mainly at NaCl concentrations of 60 mM to 100 mM. Whereas, the concentrations below and over this range either showed notable non-specific cleavage or highly reduced enzyme cleavage efficiency, respectively.

**Figure 2.**
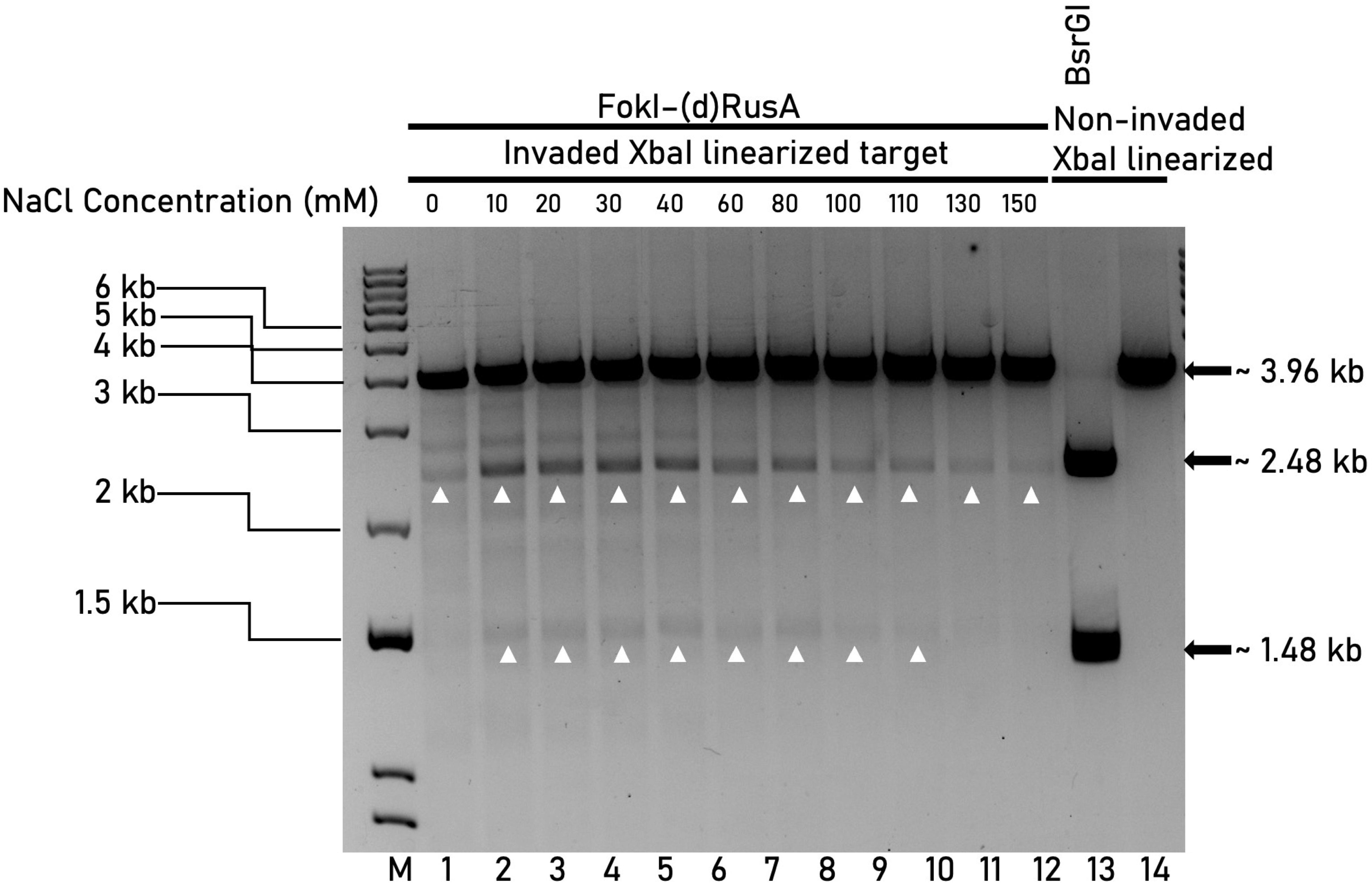
Optimization of NaCl concentrations for FokI-(d)RusA mediated target cleavage. Gel image shows the activity of FokI-(d)RusA in the presence of an increasing concentrations of NaCl on XbaI linearized pUC19-γPNA1-151 invaded with γPNA1 (Lanes 1-12). FokI-(d)RusA cleavage products are indicated by arrowheads. BsrGI and XbaI restriction enzymes cleaved and XbaI-linearized non-invaded pUC19-γPNA1-151 was included in lane 13 and 14, respectively as positive size controls. Lane M shows the 1-kb plus DNA marker.

The cleavage activity of FokI-(d)RusA was also tested on γPNA1-invaded circular pUC19-γPNA1-151 DNA using the optimal NaCl concentrations (60 mM and 100 mM) to evaluate its impact on enzyme specificity and efficiency (Figure S9). Increasing concentrations of FokI- (d)RusA (10, 20, 40, 50, 70, 90 nM) were used. The results reveal that from 60 mM NaCl to 100 mM NaCl, the cleavage efficiency was significantly compromised and resulting in very faint bands at higher protein concentrations. Whereas the non-specific cleavage was slightly reduced. Suggesting that while NaCl can stabilize specific DNA-protein interactions and reduce non-specific activity, it also lowers the overall efficiency of FokI-(d)RusA, particularly on circular DNA targets. Due to FokI-(d)RusA undesirable behavior on invaded circular targets, only invaded linear targets were used in all the optimization and investigation experiments.

### Optimization of cleavage temperature, buffer, and time

Further optimizations such as cleavage temperatures, buffer conditions and cleavage time points were necessary to generate high-quality and precise fragments for *in vitro* larger fragment cloning applications. To evaluate FokI-(d)RusA activity at different temperatures, linear pUC19-γPNA1-151 was invaded by γPNA1 at 37°C, then treated with 50 nM FokI-(d)RusA and subjected to varying increasing temperatures 4, 10, 20, 30, 37, 45, 55, 65, 75, 85, and 90°C (Figure 3). Evident cleavage was observed at temperatures ranging between 20 to 55°C, with 30°C being the optimal cleavage temperature. Temperatures below or over 30°C within the mentioned range either had clean but low cleavage activity or high cleavage activity but with significant non-specific cleavage, respectively. Based on this data, the cleavage temperature for all future experiments was set to 30°C.

**Figure 3.**
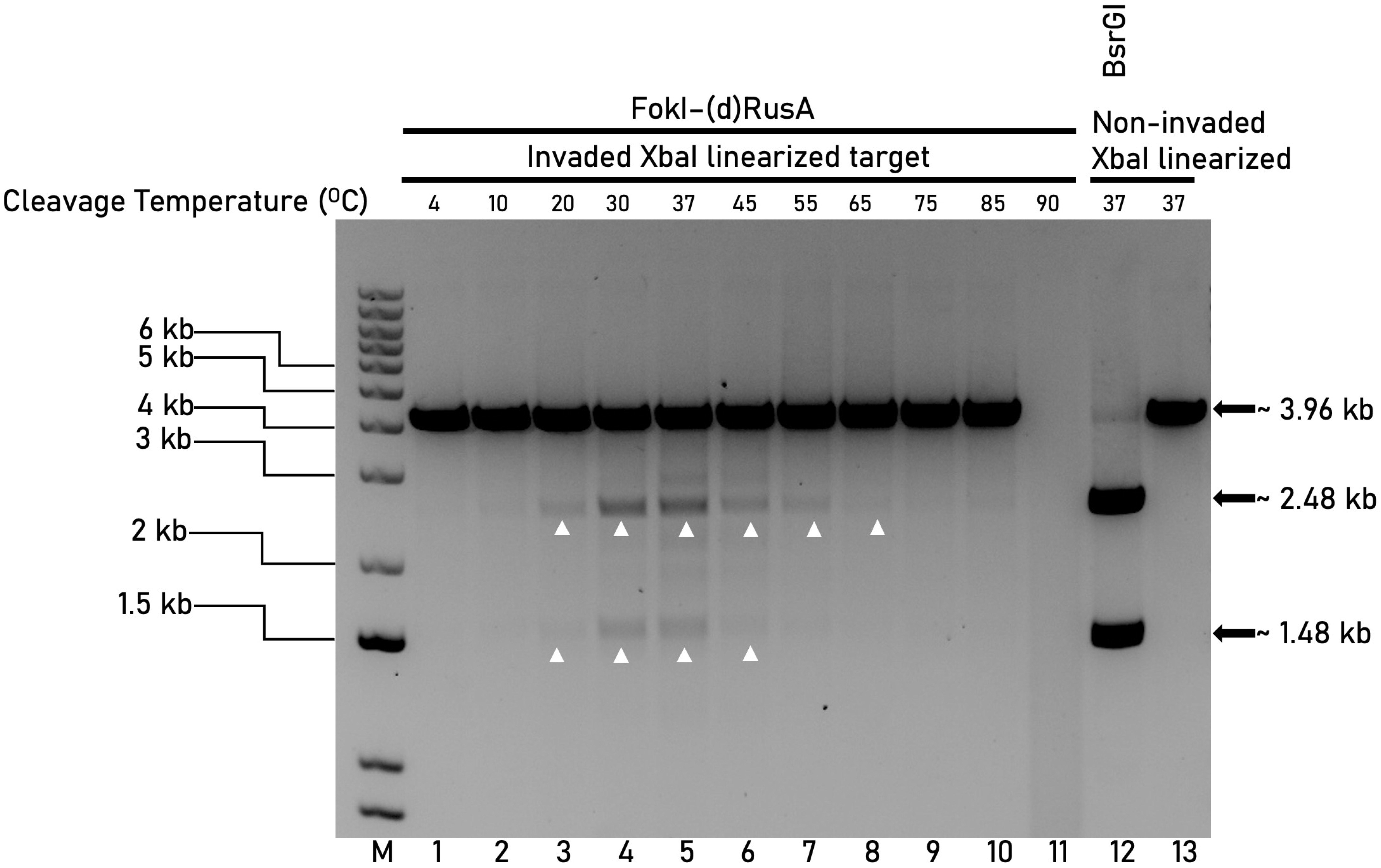
Optimization of temperature for FokI-(d)RusA mediated cleavage on invaded linear target. Gel image shows the activity of FokI-(d)RusA on XbaI linearized pUC19-γPNA1-151 invaded with γPNA1 at varying temperatures (Lanes 1-11). FokI-(d)RusA cleavage sites are indicated by arrowheads. BsrGI and XbaI restriction enzymes cleaved and XbaI-linearized non-invaded pUC19-γPNA1-151 was included in lane 12 and 13, respectively as positive size controls. Lane M shows the 1-kb plus DNA marker.

To investigate the effect of different buffers on FokI-(d)RusA and conclude on optimal buffer conditions, we tested FokI-(d)RusA cleavage activity in different buffers. We treated pUC19-γPNA1-151 invaded with γPNA1 with FokI-(d)RusA and tested in no buffer, NEB-rCutsmart, NEB-r1.1, NEB-r2.1, NEB-r3.1, HEPES, MOPS, and PBS buffer (Table S7). FokI-(d)RusA was able to induce cleavage only in NEB-rCutsmart, NEB-r2.1, and NEB-r1.1, in a decreasing cleavage strength order, while in no buffer, HEPES, MOPS, and PBS buffers, no cleavage was observed (Figure 4). NEB-rCutsmart, with a balanced ionic strength and near-neutral pH (∼7.9), was identified as the optimal buffer, providing the necessary cations (Mg²□, Na□, K□) to support specific and efficient cleavage.

**Figure 4.**
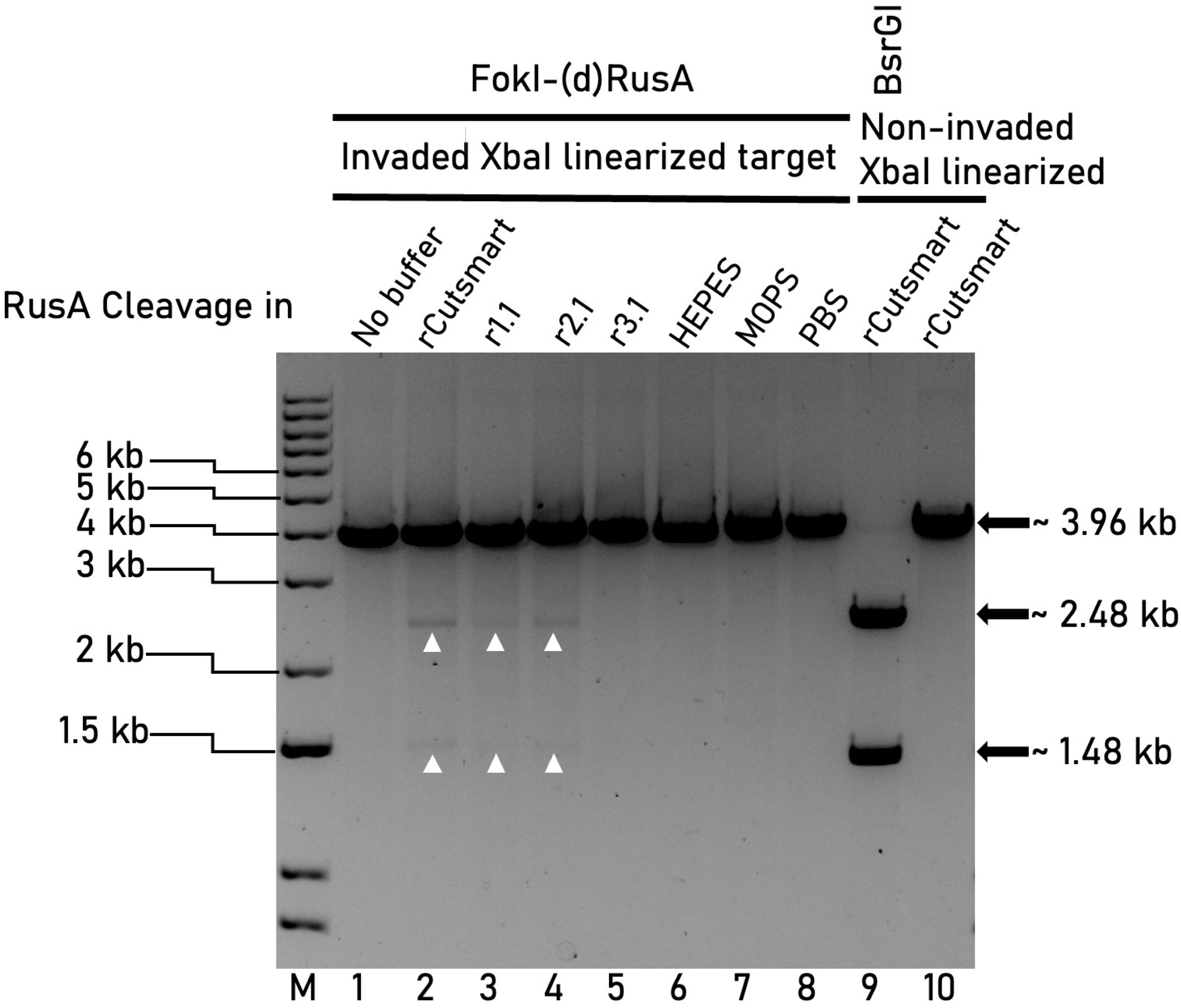
Optimization of buffer for FokI-(d)RusA mediated cleavage on invaded linear target. Gel image shows the activity of FokI-(d)RusA on XbaI linearized pUC19-γPNA1-151 invaded with γPNA1 in different buffers (Lanes 1-8). FokI-(d)RusA cleavage sites are indicated by arrowheads. BsrGI and XbaI restriction enzymes cleaved and XbaI-linearized non-invaded pUC19-γPNA1-151 was included in lane 9 and 10, respectively as positive size controls. Lane M shows the 1-kb plus DNA marker.

FokI-(d)RusA cleavage activity was also evaluated at different time-points by incubating γPNA1-invaded linear pUC19-γPNA1-151 target with FokI-(d)RusA for 0, 5, 10, 15, 20, 25, 30, 35, 40, 45, 50, 55, 60 minutes at 30°C in NEB-rCutsmart. The band release was notably visible at time 20 mins and the cleavage strength increased steadily with the increasing temperatures until 60 minutes (Figure 5). Results also confirmed that 60 minutes cleavage time is sufficient to produce observable clean fragments. In all the future cleavage experiments, the reactions were incubated for 60 minutes.

**Figure 5.**
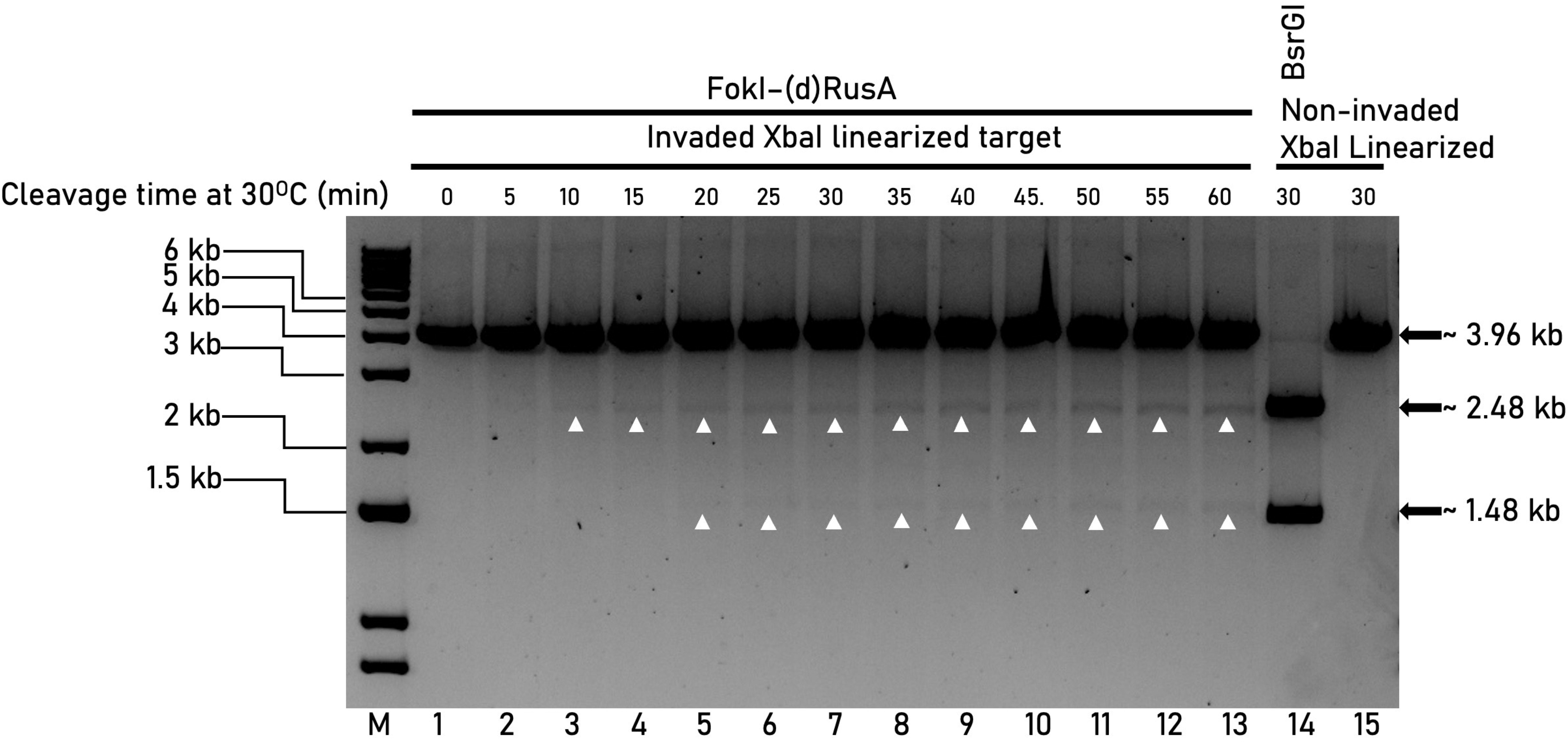
Optimization of time for FokI-(d)RusA mediated cleavage on invaded linear target. Gel image shows the activity of FokI-(d)RusA on γPNA1 invaded XbaI linearized pUC19-γPNA1-151 at different time points (Lanes 1-13). FokI-(d)RusA cleavage sites are indicated by arrowheads. BsrGI and XbaI restriction enzymes cleaved and XbaI-linearized non-invaded pUC19-γPNA1-151 was included in lane 14 and 15, respectively as positive size controls. Lane M shows the 1-kb plus DNA marker.

### Large fragment excision via PC-FIRA multiplexing activity

We assessed the PC-FIRA system capacity to manipulate large fragments for use in downstream applications such as cloning and genome assembly. A designed clone pUC19-γPNA1&1-245 with two γPNA1 target sites separated by a 1291 bp spacer was linearized with XmnI, invaded with γPNA1 (Files S2b). The invaded target was treated with FokI-(d)RusA using optimal cleavage reaction conditions. Results showed the release of 1291 bp fragment, a band of interest on the agarose gel confirming the ability of PC-FIRA system for the cleavage of larger fragment (Figure 6). Whereas the negative control samples did not show any band release confirming PC-FIRA system specificity. Different restriction enzymes cleavage of pUC19-γPNA1&1-245 plasmid showing three fragments at 1881 bp, 1314 bp, and 603 bp, as size markers for the PC-FIRA system cleavage. The 1314 bp fragment corresponds to the approximate size of the 1291 bp spacer of interest. Similar to the established PG-T7EI system, PC-FIRA was able to successfully facilitate cleavage and release of a fragment >1000 bp. Following on this, cloning of the excised fragment can be attempted.

**Figure 6.**
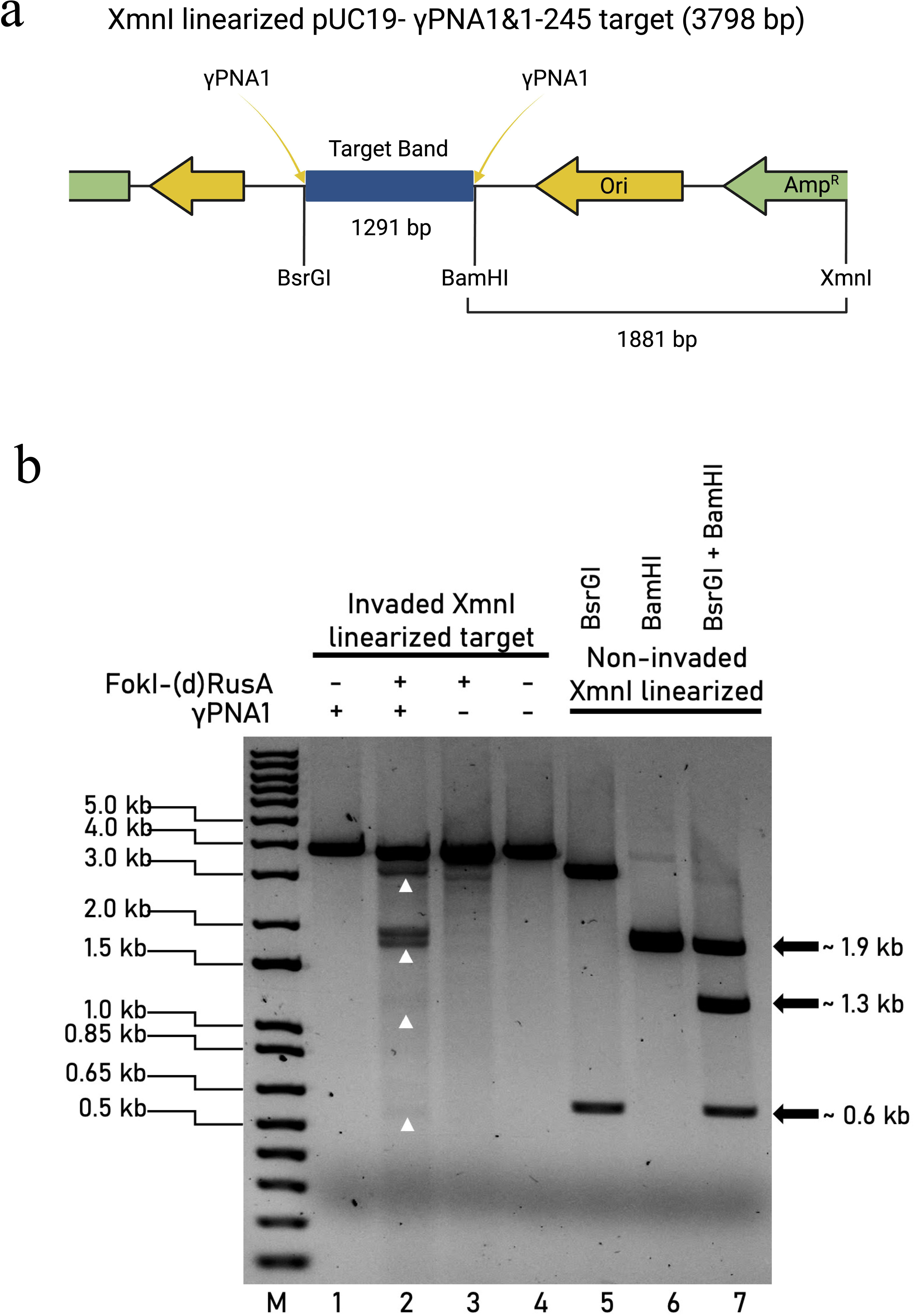
Multiplexed cleavage of invaded target using the PC-FIRA system. (a) Plasmid map for XmnI linearized pUC19-γPNA1&1-245 target. Restriction enzyme sites and γPNA1 invasion sites are indicated on the plasmid map. Size of fragment released after FokI-(d)RusA mediated cleavage are also indicated on the map. (b) Gel image showing the multiplex cleavage activity of FokI-(d)RusA on XmnI linearized pUC19-γPNA1&1-245 target, invaded with γPNA1 (Lane 2). Only γPNA1 invaded XmnI linearized pUC19-γPNA1&1-245 target (Lane 1); non-invaded XmnI linearized pUC19-γPNA1&1-245 target treated with FokI-(d)RusA (Lane 3); and XmnI linearized pUC19-γPNA1&1-245 plasmids (Lane 4), are included as negative controls. Different restriction enzyme controls, XmnI + BsrGI (Lane 5), XmnI + BamHI (Lane 6), and XmnI + BsrGI + BamHI (Lane 6) on non-invaded XbaI linearized pUC19-γPNA1&1-245 was included as positive size controls. Lane M shows the 1-kb plus DNA marker.

## CONCLUSION

The PC-FIRA system has addressed multiple barriers that limits previously established SSNs and SGNs. By removing the dependence on specific sequence motifs like PAMs, the system can target more diverse regions of the genome. Moreover, the PC-FIRA system can induce DSBs with a single PNA invasion, therefore, it offers a more cost-efficient and less complex gene editing tool. It enhances specificity by utilizing PNAs to target complex and highly specific DNA structures, reducing off-target effects.

Optimizing size is a critical factor in genome engineering applications, particularly for delivery and manipulation within cellular environments. The small size of RusA is advantageous in overcoming delivery concerns. However, the fusion of FokI to (d)RusA introduces complexities that could influence both (d)RusA and FokI activities. Considering that both FokI and RusA require dimerization for proper function, three scenarios for dimerization, binding, and cleavage activity were proposed accounting for protein conformational stability and dimerization-dependent activity (Figure S10). After successful PNA invasion (Figure S10A), three scenarios were considered, each involving different orders of dimerization and binding, accounting for factors such as protein conformational stability and dimerization-dependent catalytic activity.

The most likely scenario (Figure S10B-2-2) which should be favorable due to reduced spatial constraints on the FokI domain. The least possible scenario (Figure S10B-1) involves the dimerization of two RusA and two FokI domains between two free FokI-(d)RusA monomers, which then bind to PNA-invaded site and induces cleavage. However, this scenario is unlikely, as the catalytic activity of FokI-(d)RusA depends on the correct positioning of target within its FokI domain *^35^*which is mediated through RusA binding activity. In order to confirm such theories and provide more insights into this system, the use of imaging techniques, such as Cryo-EM and X-ray crystallography, can be utilized. Studies have shown that careful design and proper understanding of the dimerization activity in FokI-based nucleases, is crucial to minimize toxicity and off-target effects*^36, 37^*. Understanding these dynamics is crucial for designing a valuable tool for further genome engineering applications.

The presented approach leverages FokI, RusA, and PNAs to construct an efficient and accurate gene-editing system. The high specificity, considerable in vitro efficiency, and easy seamless customizability of the PC-FIRA system makes it a viable tool that warrants further exploration to achieve its full potential. Through this work, we identified an ultra-compact resolvase with potential in DNA manipulation applications. Future research should be focused on investigating the PC-FIRA system within different genomic contexts, potentially making it suitable for diverse applications in genome editing, therapeutic development, and synthetic biology.

## Author contributions

A.S, G.S.R and M.M designed the research; A.S, G.S.R, and Q.W performed the experiments; A.S, G.S.R and M.M analyzed data and wrote the paper.

## ACKNOWLEDGEMENTS

We would like to thank members of the genome engineering and synthetic biology laboratory for their discussions and help.

## REFERENCES

[1] Urnov, F. D., Rebar, E. J., Holmes, M. C., Zhang, H. S., and Gregory, P. D. (2010) Genome editing with engineered zinc finger nucleases, Nature Reviews Genetics 11, 636–646.

[2] Jinek, M., Chylinski, K., Fonfara, I., Hauer, M., Doudna, J. A., and Charpentier, E. (2012) A programmable dual-RNA-guided DNA endonuclease in adaptive bacterial immunity, Science 337, 816–821.

[3] Joung, J. K., and Sander, J. D. (2013) TALENs: a widely applicable technology for targeted genome editing, Nature reviews Molecular cell biology 14, 49–55.

[4] Hillary, V. E., and Ceasar, S. A. (2023) A Review on the Mechanism and Applications of CRISPR/Cas9/Cas12/Cas13/Cas14 Proteins Utilized for Genome Engineering, Molecular Biotechnology 65, 311–325.

[5] Tyumentseva, M., Tyumentsev, A., and Akimkin, V. (2023) CRISPR/Cas9 Landscape: Current State and Future Perspectives, International Journal of Molecular Sciences 24, 16077.

[6] Cui, Y., Xu, J., Cheng, M., Liao, X., and Peng, S. (2018) Review of CRISPR/Cas9 sgRNA design tools, Interdisciplinary Sciences: Computational Life Sciences 10, 455–465.

[7] Taha, E. A., Lee, J., and Hotta, A. (2022) Delivery of CRISPR-Cas tools for in vivo genome editing therapy: Trends and challenges, Journal of Controlled Release 342, 345–361.

[8] Hutvagner, G., and Simard, M. J. (2008) Argonaute proteins: key players in RNA silencing, Nature Reviews Molecular Cell Biology 9, 22–32.

[9] Marsic, T., Gundra, S. R., Wang, Q., Aman, R., Mahas, A., and Mahfouz, Magdy M. (2023) Programmable site-specific DNA double-strand breaks via PNA-assisted prokaryotic Argonautes, Nucleic Acids Research 51, 9491–9506.

[10] Nielsen, P. E., Egholm, M., Berg, R. H., and Buchardt, O. (1991) Sequence-selective recognition of DNA by strand displacement with a thymine-substituted polyamide, Science 254, 1497–1500.

[11] Singh, K. R., Sridevi, P., and Singh, R. P. (2020) Potential applications of peptide nucleic acid in biomedical domain, Engineering Reports 2, e12238.

[12] Bahal, R., Ali McNeer, N., Quijano, E., Liu, Y., Sulkowski, P., Turchick, A., Lu, Y.-C., Bhunia, D. C., Manna, A., Greiner, D. L., Brehm, M. A., Cheng, C. J., López-Giráldez, F., Ricciardi, A., Beloor, J., Krause, D. S., Kumar, P., Gallagher, P. G., Braddock, D. T., Mark Saltzman, W., Ly, D. H., and Glazer, P. M. (2016) In vivo correction of anaemia in β-thalassemic mice by γPNA-mediated gene editing with nanoparticle delivery, Nature Communications 7, 13304.

[13] Jiang, W., Aman, R., Ali, Z., Rao, G. S., and Mahfouz, M. (2023) PNA-pdx: versatile peptide nucleic acid-based detection of nucleic acids and SNPs, Analytical Chemistry 95, 14209–14218.

[14] Branzei, D., and Foiani, M. (2008) Regulation of DNA repair throughout the cell cycle, Nature Reviews Molecular Cell Biology 9, 297–308.

[15] Wyatt, H. D., and West, S. C. (2014) Holliday junction resolvases, Cold Spring Harb Perspect Biol 6, a023192.

[16] Hadden, J. M., Convery, M. A., Déclais, A.-C., Lilley, D. M., and Phillips, S. E. (2001) Crystal structure of the Holliday junction resolving enzyme T7 endonuclease I, nature structural biology 8, 62–67.

[17] Kobayashi, Y., Odahara, M., Sekine, Y., Hamaji, T., Fujiwara, S., Nishimura, Y., and Miyagishima, S.-y. (2020) Holliday Junction Resolvase MOC1 Maintains Plastid and Mitochondrial Genome Integrity in Algae and Bryophytes, Plant Physiology 184, 1870–1883.

[18] Aman, R., Syed, M. M., Saleh, A., Melliti, F., Gundra, Sivakrishna R., Wang, Q., Marsic, T., Mahas, A., and Mahfouz, Magdy M. (2024) Peptide nucleic acid-assisted generation of targeted double-stranded DNA breaks with T7 endonuclease I, Nucleic Acids Research 52, 3469–3482.

[19] Sivakrishna Rao, G., Saleh, A. H., Melliti, F., Muntjeeb, S., and Mahfouz, M. (2024) Harnessing Peptide Nucleic Acids and the Eukaryotic Resolvase MOC1 for Programmable, Precise Generation of Double-Strand DNA Breaks, Anal Chem 96, 2599–2609.

[20] San-Segundo, P. A., and Clemente-Blanco, A. (2020) Resolvases, dissolvases, and helicases in homologous recombination: clearing the road for chromosome segregation, Genes 11, 71.

[21] Chan, S. N., Vincent, S. D., and Lloyd, R. G. (1998) Recognition and manipulation of branched DNA by the RusA Holliday junction resolvase of Escherichia coli, Nucleic Acids Research 26, 1560–1566.

[22] Sharples, G. J., Chan, S. N., Mahdi, A. A., Whitby, M. C., and Lloyd, R. G. (1994) Processing of intermediates in recombination and DNA repair: identification of a new endonuclease that specifically cleaves Holliday junctions, The EMBO Journal 13, 6133–6142.

[23] West, S. C. (1997) Processing of recombination intermediates by the RuvABC proteins, Annu Rev Genet 31, 213-244.

[24] Macmaster, R., Sedelnikova, S., Baker, P. J., Bolt, E. L., Lloyd, R. G., and Rafferty, J. B. (2006) RusA Holliday junction resolvase: DNA complex structure—insights into selectivity and specificity, Nucleic Acids Research 34, 5577–5584.

[25] Boddy, M. N., Gaillard, P. H. L., McDonald, W. H., Shanahan, P., Yates, J. R., 3rd, and Russell, P. (2001) Mus81-Eme1 are essential components of a Holliday junction resolvase, Cell 107, 537-548.

[26] Doe, C. L., Ahn, J. S., Dixon, J., and Whitby, M. C. (2002) Mus81-Eme1 and Rqh1 involvement in processing stalled and collapsed replication forks, J Biol Chem 277, 32753–32759.

[27] Pernstich, C., and Halford, S. E. (2012) Illuminating the reaction pathway of the FokI restriction endonuclease by fluorescence resonance energy transfer, Nucleic acids research 40, 1203–1213.

[28] Guilinger, J. P., Thompson, D. B., and Liu, D. R. (2014) Fusion of catalytically inactive Cas9 to FokI nuclease improves the specificity of genome modification, Nature Biotechnology 32, 577–582.

[29] Wang, Q., Rao, G. S., Marsic, T., Aman, R., and Mahfouz, M. (2024) Fusion of FokI and catalytically inactive prokaryotic Argonautes enables site-specific programmable DNA cleavage, J Biol Chem 300, 107720.

[30] Abramson, J., Adler, J., Dunger, J., Evans, R., Green, T., Pritzel, A., Ronneberger, O., Willmore, L., Ballard, A. J., Bambrick, J., Bodenstein, S. W., Evans, D. A., Hung, C.-C., O’Neill, M., Reiman, D., Tunyasuvunakool, K., Wu, Z., Žemgulytė, A., Arvaniti, E., Beattie, C., Bertolli, O., Bridgland, A., Cherepanov, A., Congreve, M., Cowen-Rivers, A. I., Cowie, A., Figurnov, M., Fuchs, F. B., Gladman, H., Jain, R., Khan, Y. A., Low, C. M. R., Perlin, K., Potapenko, A., Savy, P., Singh, S., Stecula, A., Thillaisundaram, A., Tong, C., Yakneen, S., Zhong, E. D., Zielinski, M., Žídek, A., Bapst, V., Kohli, P., Jaderberg, M., Hassabis, D., and Jumper, J. M. (2024) Accurate structure prediction of biomolecular interactions with AlphaFold 3, Nature 630, 493–500.

[31] Bolt, E. L., and Lloyd, R. G. (2002) Substrate specificity of RusA resolvase reveals the DNA structures targeted by RuvAB and RecG in vivo, Mol Cell 10, 187–198.

[32] Ramirez, C. L., and Joung, J. K. (2013) Engineered Zinc Finger Nucleases for Targeted Genome Editing, In Site-directed insertion of transgenes (Renault, S., and Duchateau, P., Eds.), pp 121-145, Springer Netherlands, Dordrecht.

[33] Scott, S., Xu, Zhi M., Kouzine, F., Berard, D. J., Shaheen, C., Gravel, B., Saunders, L., Hofkirchner, A., Leroux, C., Laurin, J., Levens, D., Benham, C. J., and Leslie, S. R. (2018) Visualizing structure-mediated interactions in supercoiled DNA molecules, Nucleic Acids Research 46, 4622–4631.

[34] Kumar, A., Daripa, P., Rasool, K., Chakraborty, D., Jain, N., and Maiti, S. (2024) Deciphering the Thermodynamic Landscape of CRISPR/Cas9: Insights into Enhancing Gene Editing Precision and Efficiency, The Journal of Physical Chemistry B 128, 8409–8422.

[35] Sanders, K. L., Catto, L. E., Bellamy, S. R. W., and Halford, S. E. (2009) Targeting individual subunits of the FokI restriction endonuclease to specific DNA strands, Nucleic Acids Research 37, 2105–2115.

[36] Szczepek, M., Brondani, V., Büchel, J., Serrano, L., Segal, D. J., and Cathomen, T. (2007) Structure-based redesign of the dimerization interface reduces the toxicity of zinc-finger nucleases, Nature Biotechnology 25, 786–793.

[37] Tsai, S. Q., Wyvekens, N., Khayter, C., Foden, J. A., Thapar, V., Reyon, D., Goodwin, M. J., Aryee, M. J., and Joung, J. K. (2014) Dimeric CRISPR RNA-guided FokI nucleases for highly specific genome editing, Nature Biotechnology 32, 569–576.

