## Supplementary information for "Fusions of catalytically inactive RusA to FokI nuclease coupled with PNA enable programable site-specific double-stranded DNA breaks"

**Supplementary figures**

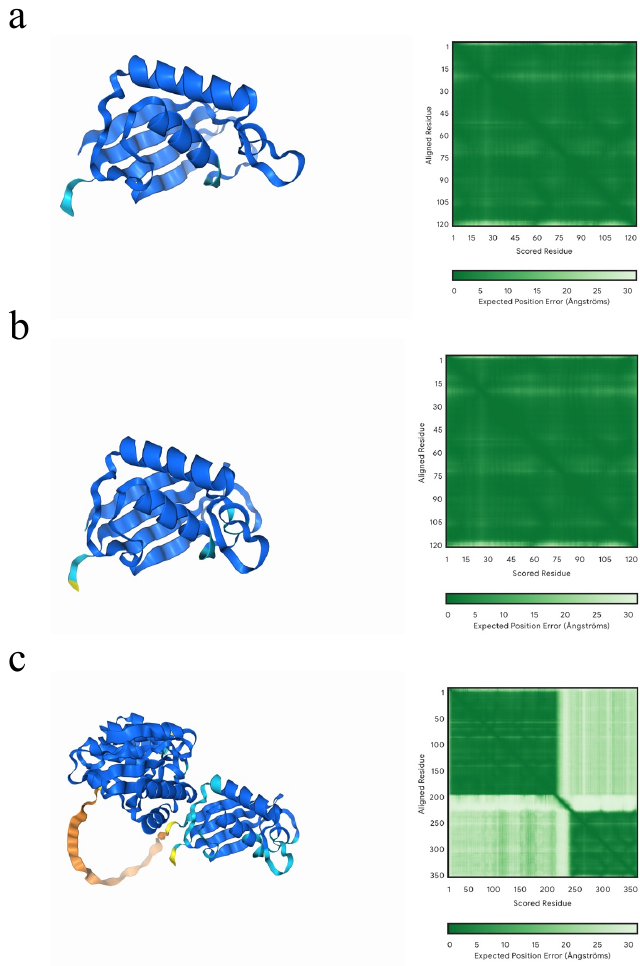

**Figure S1.** Predicted structures of (wt)RusA, (d)RusA and FokI-(d)RusA using AlphaFold 3.0. The α/β fold models of (a) (wt)RusA, (b) (d)RusA, and (c) FokI-(d)RusA fusion. Structural confidence is indicated by color and are based on the (pLDDT) score, which ranges from 0 to 100. Scores above 90 (blue) indicate very high confidence, 70–90 (cyan) moderate confidence, 50–70 (yellow) low confidence, and below 50 (orange) very low confidence. Predicted Position Error (PPE) graph is also presented for each protein. Lower errors (closer to 0 Å) are indicated in green, while higher values (closer to 30 Å) are indicated in white.

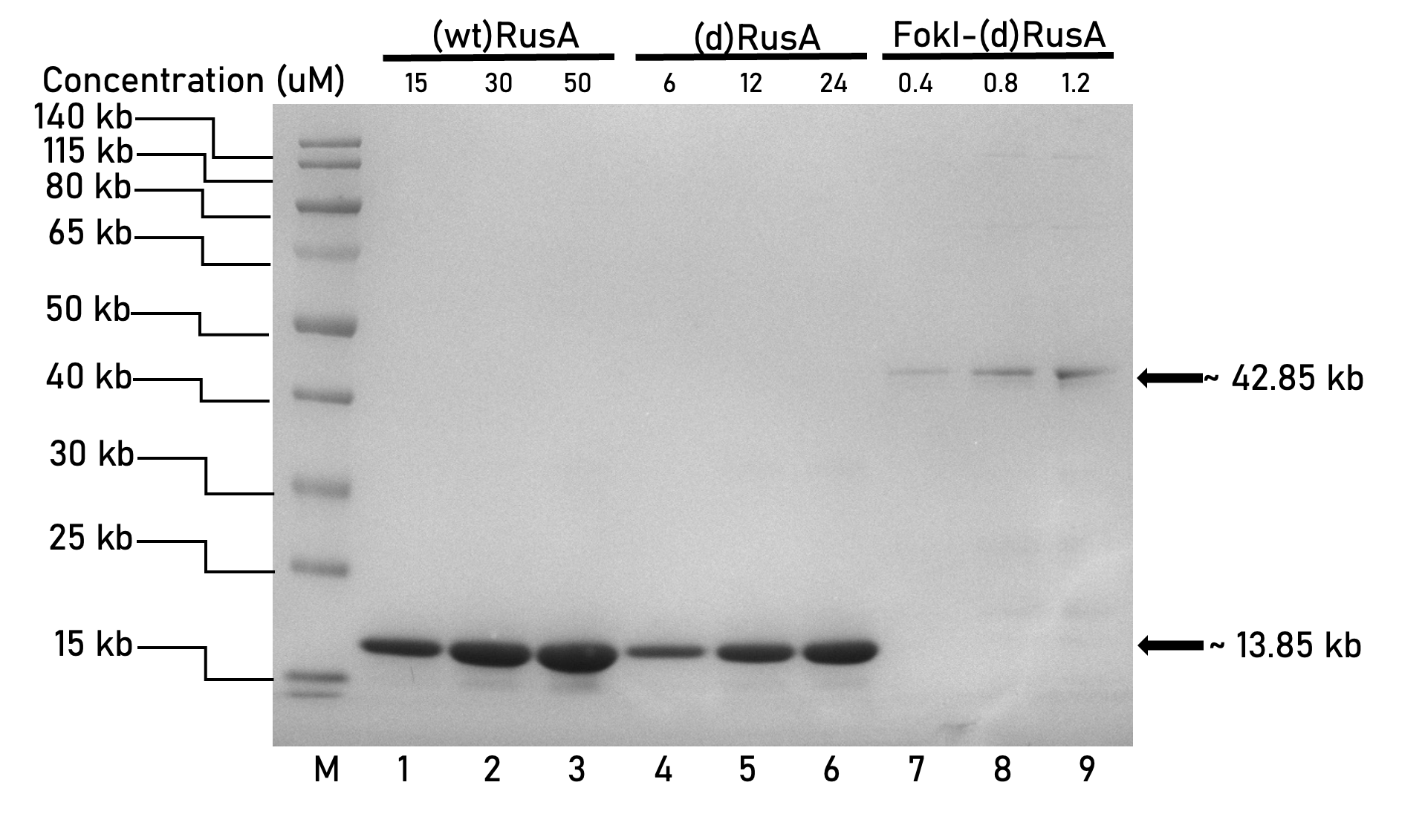

**Figure S2.** SDS-PAGE analysis of (wt)RusA, (d)RusA, and FokI-(d)RusA. Proteins were analyzed by SDS-PAGE to qualitatively evaluate purity. Increasing concentrations of each protein were loaded onto the gel with (wt)RusA (Lane 1-3), (d)RusA (Lane 4-6), and FokI-(d)RusA (Lane 7-8). Lane M is the PageRuler^TM^ prestained protein ladder.

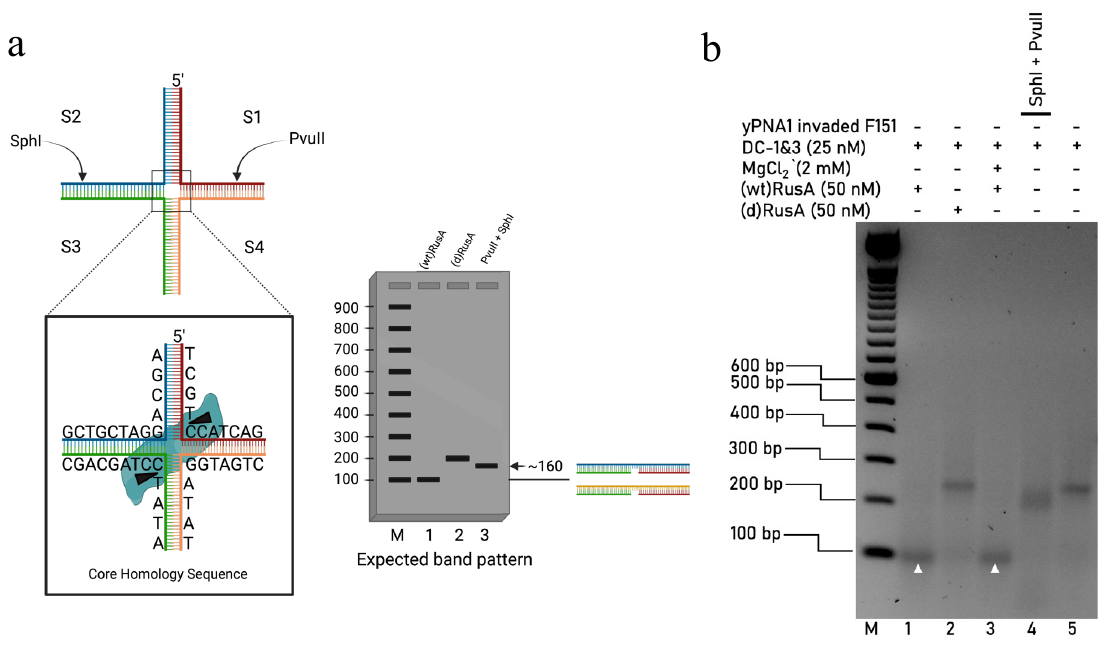

**Figure S3.** Design of DC-1&3 HJ and its expected and observed cleavage pattern by (wt)RusA and (d)RusA. **(**a) Design of DC-1&3 HJs with approximate cutting sites for both PuvII and SphI used indicated by arrows; a zoom in on the adopted core homology sequences with cutting sites for (wt)RusA indicated by arrowheads; and Expected band pattern for (wt)RusA and (d)RusA activity on DC-1&3, and PvuII and SphI as restriction control. (b) Gel image shows the activity of (wt)RusA and (d)RusA on DC-1&3 (Lane 1-3). Restriction enzyme control on DC-1&3 was included in lanes 4. Top labelling indicates reagents, concentrations, and presence within samples. Lane M shows the 100-bp DNA marker.

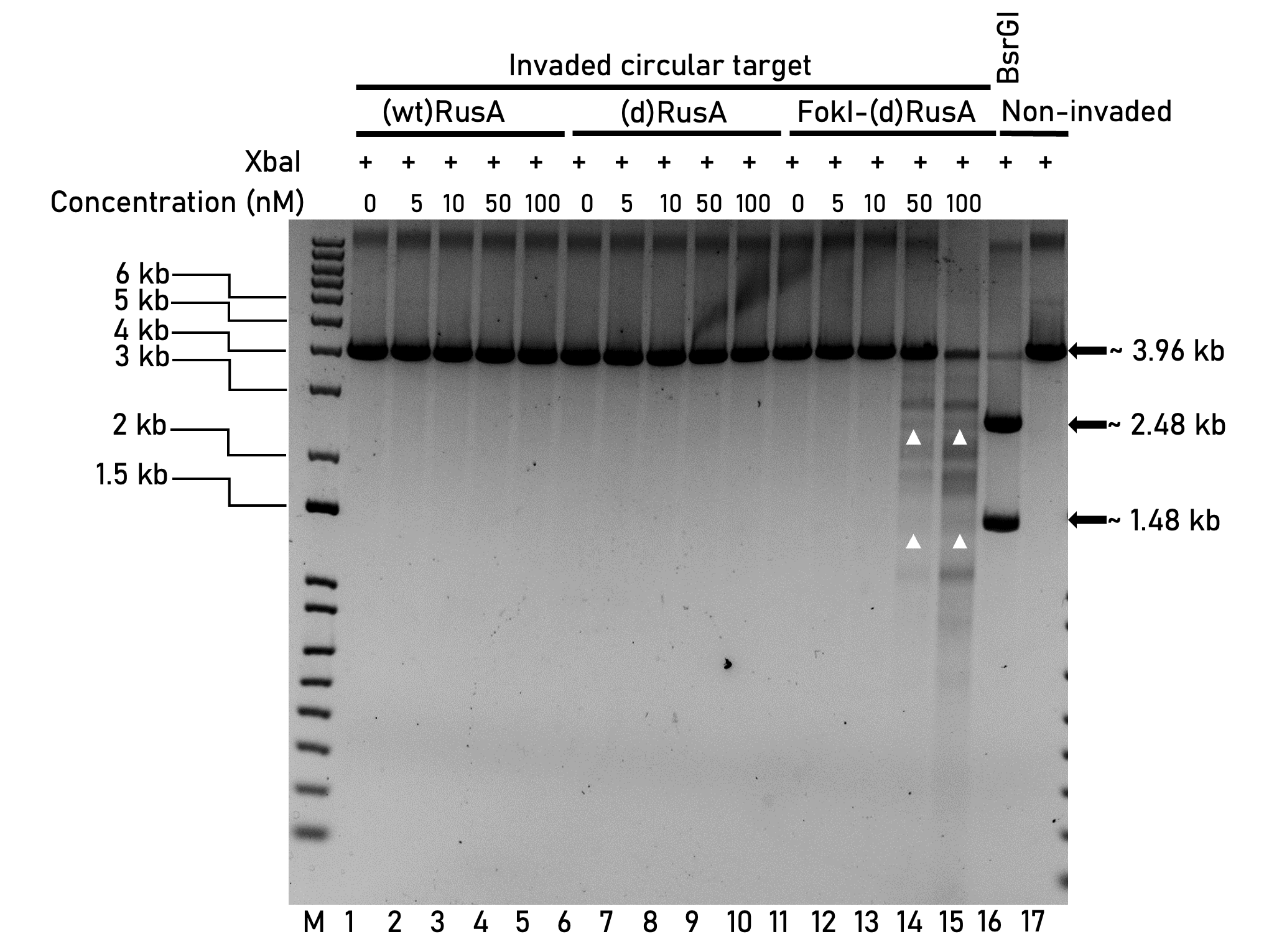

**Figure S4.** Cleavage activity of (wt)RusA, (d)RusA, and FokI-(d)RusA under optimal reaction conditions on PNA invaded circular target.  Gel image shows cleavage activity of (wt)RusA, (d)RusA, and FokI-(d)RusA at increasing concentrations (Lanes 1-15) on circular pUC19-γPNA1-151 invaded with γPNA1. (wt)RusA, (d)RusA, and FokI-(d)RusA treated samples are found in lanes (1-5), (6-10), and (11-15), respectively. XbaI was added to all samples for fragment of interest release after (wt)RusA and (d)RusA mediated cleavage. FokI-(d)RusA cleavage sites are indicated by arrowheads. Restriction enzyme control on non-invaded pUC19- γPNA1-151 was included in lane 16. Lane M shows the 1-kb plus DNA marker.

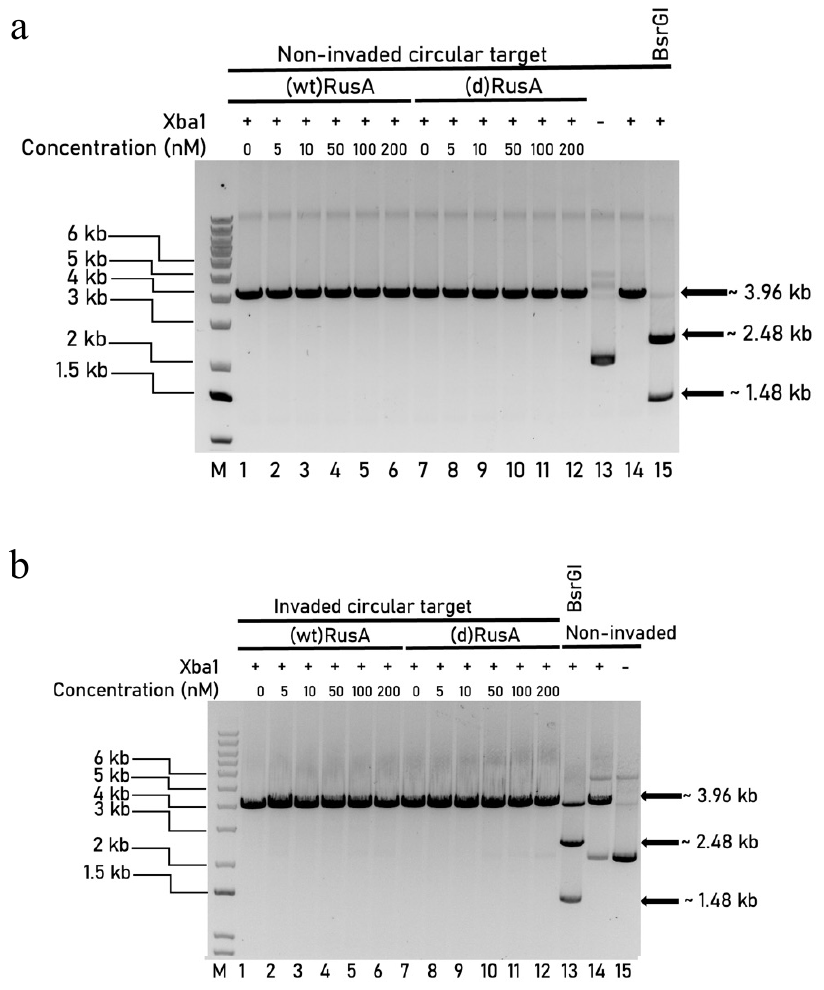

**Figure S5.** Cleavage activity of (wt)RusA and (d)RusA on non-invaded and invaded circular targets. (a) Gel image shows the activity of (wt)RusA and (d)RusA on circular pUC19-γPNA1-151 non-invaded target, with (wt)RusA and (d)RusA treated samples in lanes (1-6) and (7-12), respectively. XbaI was added to all samples for fragment of interest release after (wt)RusA and (d)RusA mediated cleavage. (b) Gel image shows the activity of (wt)RusA and (d)RusA on circular pUC19-γPNA1-151 invaded target, with (wt)RusA and (d)RusA treated samples in lanes (1-6) and (7-12), respectively. Restriction enzyme controls was included in lane 14, and 15 in both gels. Restriction enzyme controls on non-invaded circular pUC19- γPNA1-151 was included in lane 13. Lane M shows the 1-kb plus DNA marker.

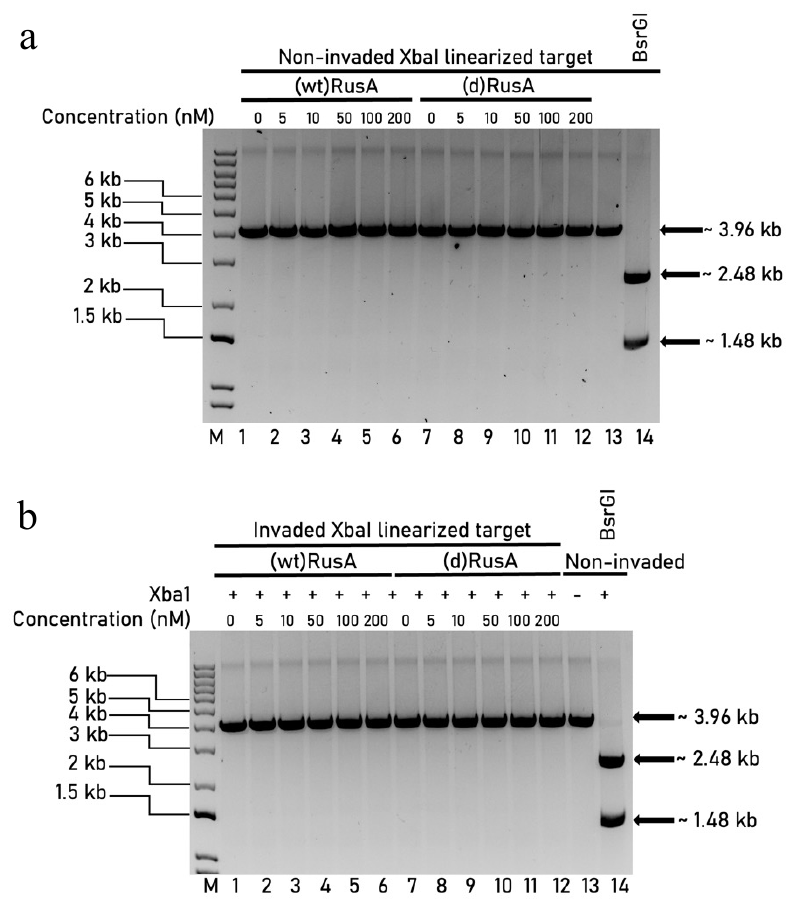

**Figure S6.** Cleavage activity of (wt)RusA (d)RusA on non-invdaded and invaded linear targets. (a) Gel image shows the activity of (wt)RusA and (d)RusA on XbaI linearized pUC19-γPNA1-151 non-invaded target, with (wt)RusA and (d)RusA treated samples in lanes (1-6) and (7-12), respectively. (b) Gel image shows the activity of (wt)RusA and (d)RusA on on XbaI linearized pUC19-γPNA1-151 invaded target, with (wt)RusA and (d)RusA treated samples in lanes (1-6) and (7-12), respectively. Restriction enzyme controls was included in lane 13 and 14. Lane M shows the 1-kb plus DNA marker.

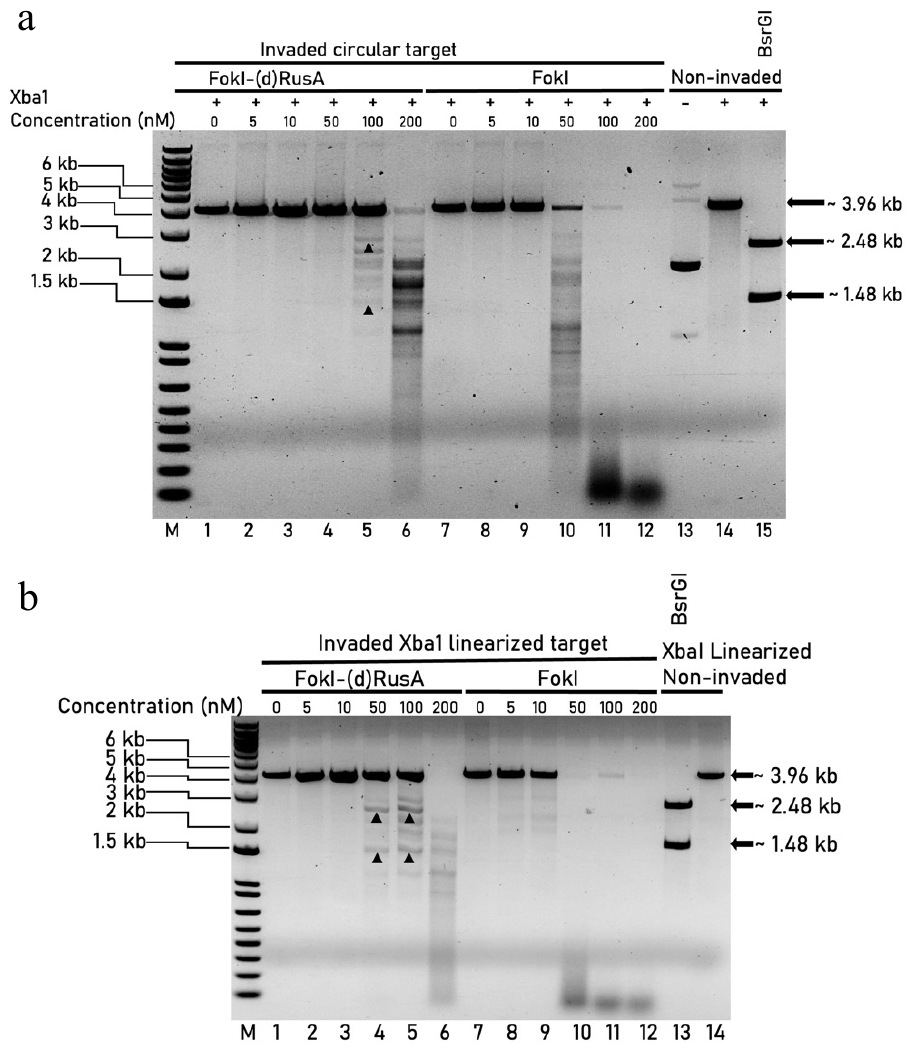

**Figure S7.** Cleavage activity of FokI-(d)RusA and FokI on invaded circular and linear targets. (a) Gel image shows the activity of FokI-(d)RusA and FokI on circular pUC19-γPNA1-151 invaded with γPNA1, with FokI-(d)RusA and FokI treated samples in lanes (1-6) and (7-12), respectively. XbaI was added to all samples for fragment of interest release after FokI-(d)RusA mediated cleavage. (b) Gel image shows the activity of FokI-(d)RusA and FokI on XbaI linearized pUC19-γPNA1-151 invaded with γPNA1, with FokI-(d)RusA and FokI treated samples in lanes (1-6) and (7-12), respectively. FokI-(d)RusA cleavage sites are indicated by arrowheads. Restriction enzyme controls on non-invaded circular and linear pUC19- γPNA1-151 was included (Lanes 14 and 15 in gel (a), and Lanes 13 and 14 in gel (b)). Lane M shows the 1-kb plus DNA marker.

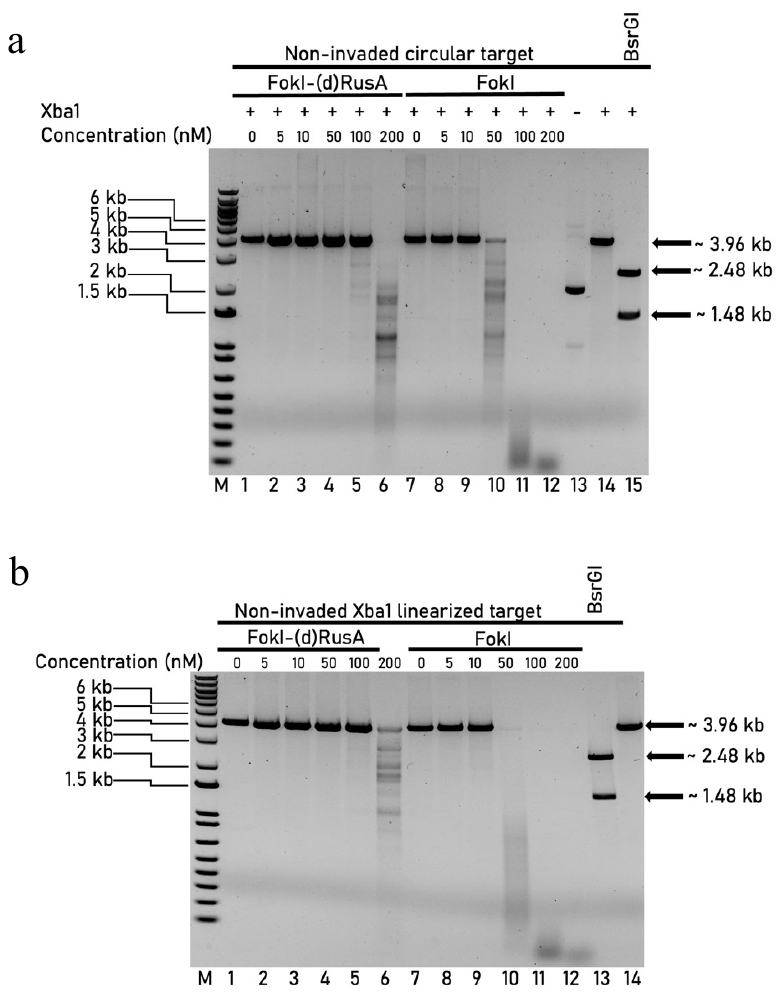

**Figure S8.** Cleavage activity of FokI-(d)RusA and FokI on non-invaded circular and linear targets. (a) Gel image shows the activity of FokI-(d)RusA and FokI on circular pUC19-γPNA1-151 non-invaded, with FokI-(d)RusA and FokI treated samples are found in lanes (1-6) and (7-12), respectively. XbaI was added to all samples for fragment of interest release after FokI-(d)RusA mediated cleavage. (b) Gel image shows the activity of FokI-(d)RusA and FokI on XbaI linearized pUC19-γPNA1-151 non-invaded target. FokI-(d)RusA and FokI treated samples are found in lanes (1-6) and (7-12), respectively. Restriction enzyme controls was included (Lanes 14 and 15 in gel (a), and Lanes 13 and 14 in gel (b)). Lane M shows the 1-kb plus DNA marker.

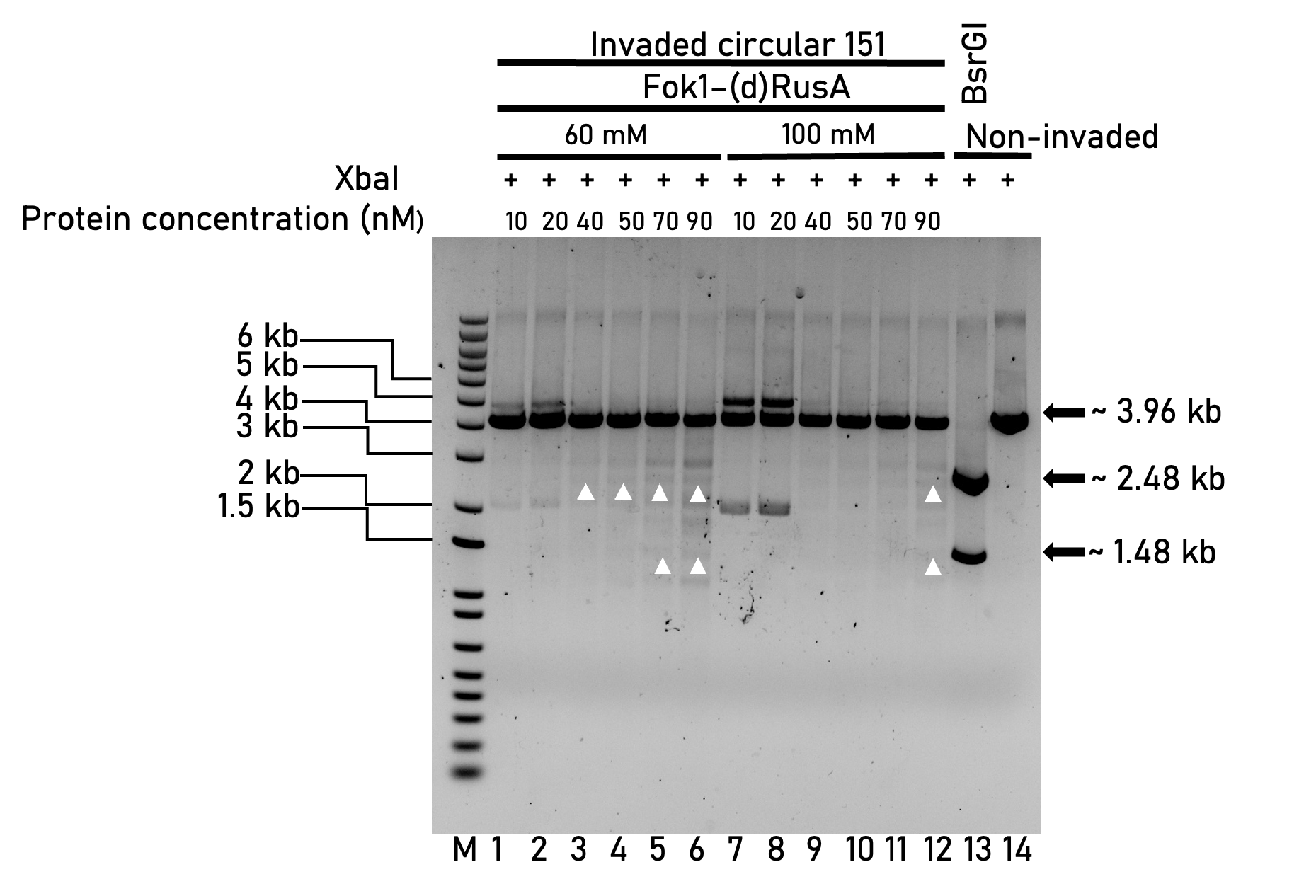

**Figure S9.** Optimization of NaCl concentrations for FokI-(d)RusA mediated cleavage on invaded circular target. Gel image shows the activity of FokI-(d)RusA at an increasing concentrations in the presence of two optimal NaCl concentrations on circular pUC19-γPNA1-151 invaded with γPNA1 (Lanes 1-12). FokI-(d)RusA cleavage sites are indicated by arrowheads. Restriction enzyme control on non-invaded circular pUC19- γPNA1-151 was included in lane 13. Lane M shows the 1-kb plus DNA marker.

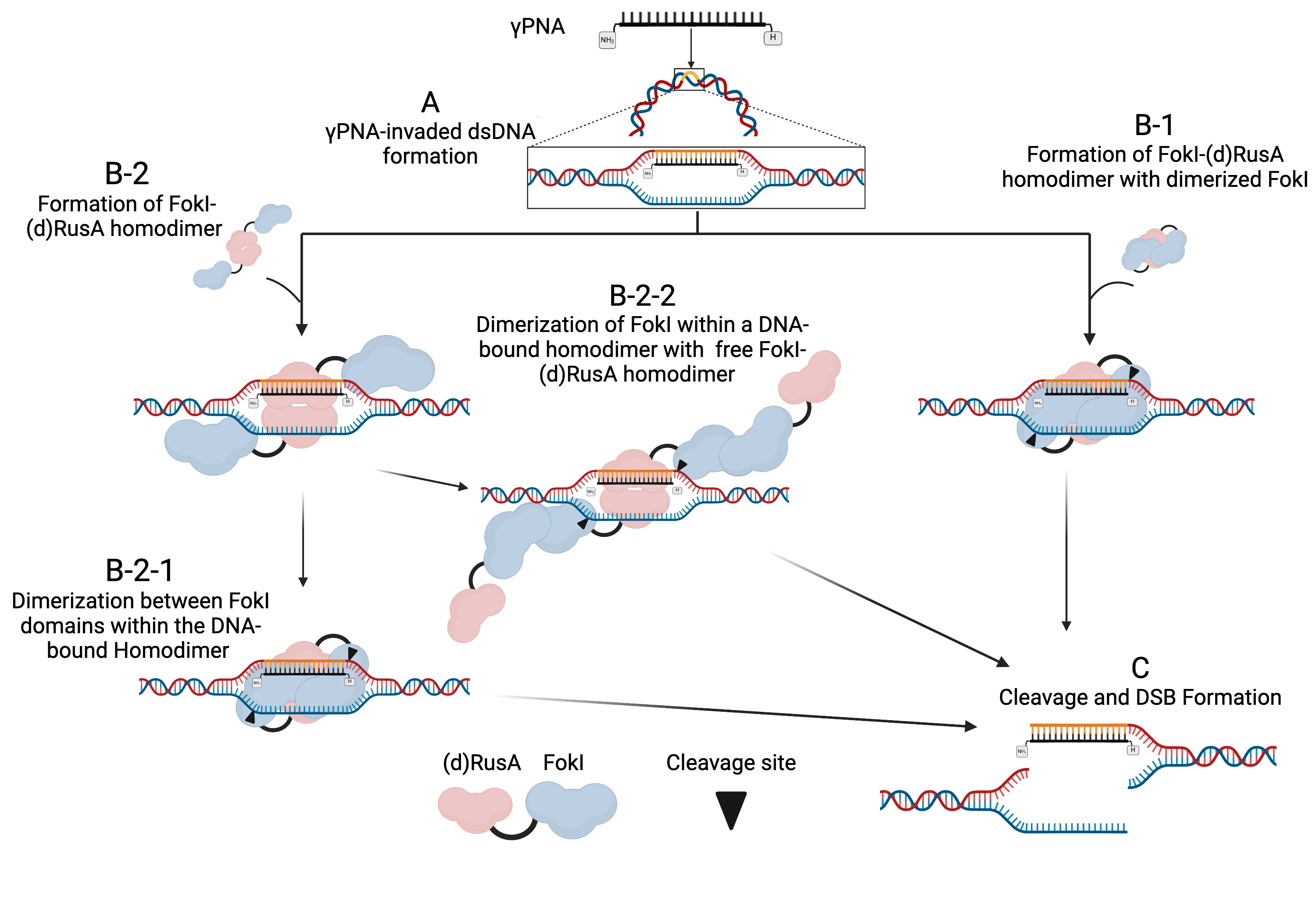

**Figure S10.** Sketch for the hypothesized three possible scenarios for the dimerization, binding, and cleavage activities of FokI-(d)RusA on γPNA-invaded DNA targets. (A) γPNA invasion of circular or linear dsDNA, forming an HJ analog. (B-1) Dimerization of FokI and (d)RusA domains then binding as a FokI-(d)RusA homodimer to γPNA-nvaded target sites — inconsistent with FokI activity and high spatial constraints. (B-2) Dimerization of (d)RusA domains then binding as a FokI-(d)RusA homodimer to γPNA-nvaded target sites (B-2-1) Dimerization of FokI domains in bound FokI-(d)RusA homodimer —high spatial constraints. (B-2-2) Dimerization of FokI domains in bound FokI-(d)RusA homodimer separately to free FokI-(d)RusA monomers — low spatial constraints. (C) Cleavage and DSBs formation. FokI and (d)RusA domains are denoted. Cleavage site of FokI-(d)RusA is indicated with arrowheads.

**Supplementary files**

1. **Protein expression plasmids used in this study**

pET28A (BamHI and XhoI are highlighted) (5379 bp)

agcgcctgatgcggtattttctccttacgcatctgtgcggtatttcacaccgCAATGGTGCACTCTCAGTACAATCTGCTCTGATGCCGCATAGTTAAGCCAGTATACACTCCGCTATCGCTACGTGACTGGGTCATGGCTGCGCCCCGACACCCGCCAACACCCGCTGACGCGCCCTGACGGGCTTGTCTGCTCCCGGCATCCGCTTACAGACAAGCTGTGACCGTCTCCGGGAGCTGCATGTGTCAGAGGTTTTCACCGTCATCACCGAAACGCGCGAGGCAGCTGCGGTAAAGCTCATCAGCGTGGTCGTGAAGCGATTCACAGATGTCTGCCTGTTCATCCGCGTCCAGCTCGTTGAGTTTCTCCAGAAGCGTTAATGTCTGGCTTCTGATAAAGCGGGCCATGTTAAGGGCGGTTTTTTCCTGTTTGGTCACTGATGCCTCCGTGTAAGGGGGATTTCTGTTCATGGGGGTAATGATACCGATGAAACGAGAGAGGATGCTCACGATACGGGTTACTGATGATGAACATGCCCGGTTACTGGAACGTTGTGAGGGTAAACAACTGGCGGTATGGATGCGGCGGGACCAGAGAAAAATCACTCAGGGTCAATGCCAGCGCTTCGTTAATACAGATGTAGGTGTTCCACAGGGTAGCCAGCAGCATCCTGCGATGCAGATCCGGAACATAATGGTGCAGGGCGCTGACTTCCGCGTTTCCAGACTTTACGAAACACGGAAACCGAAGACCATTCATGTTGTTGCTCAGGTCGCAGACGTTTTGCAGCAGCAGTCGCTTCACGTTCGCTCGCGTATCGGTGATTCATTCTGCTAACCAGTAAGGCAACCCCGCCAGCCTAGCCGGGTCCTCAACGACAGGAGCACGATCATGCGCACCCGTGGGGCCGCCATGCCGGCGATAATGGCCTGCTTCTCGCCGAAACGTTTGGTGGCGGGACCAGTGACGAAGGCTTGAGCGAGGGCGTGCAAGATTCCGAATACCGCAAGCGACAGGCCGATCATCGTCGCGCTCCAGCGAAAGCGGTCCTCGCCGAAAATGACCCAGAGCGCTGCCGGCACCTGTCCTACGAGTTGCATGATAAAGAAGACAGTCATAAGTGCGGCGACGATAGTCATGCCCCGCGCCCACCGGAAGGAGCTGACTGGGTTGAAGGCTCTCAAGGGCATCGGTCGAGATCCCGGTGCCTAATGAGTGAGCTAACTTACATTAATTGCGTTGCGCTCACTGCCCGCTTTCCAGTCGGGAAACCTGTCGTGCCAGCTGCATTAATGAATCGGCCAACGCGCGGGGAGAGGCGGTTTGCGTATTGGGCGCCAGGGTGGTTTTTCTTTTCACCAGTGAGACGGGCAACAGCTGATTGCCCTTCACCGCCTGGCCCTGAGAGAGTTGCAGCAAGCGGTCCACGCTGGTTTGCCCCAGCAGGCGAAAATCCTGTTTGATGGTGGTTAACGGCGGGATATAACATGAGCTGTCTTCGGTATCGTCGTATCCCACTACCGAGATATCCGCACCAACGCGCAGCCCGGACTCGGTAATGGCGCGCATTGCGCCCAGCGCCATCTGATCGTTGGCAACCAGCATCGCAGTGGGAACGATGCCCTCATTCAGCATTTGCATGGTTTGTTGAAAACCGGACATGGCACTCCAGTCGCCTTCCCGTTCCGCTATCGGCTGAATTTGATTGCGAGTGAGATATTTATGCCAGCCAGCCAGACGCAGACGCGCCGAGACAGAACTTAATGGGCCCGCTAACAGCGCGATTTGCTGGTGACCCAATGCGACCAGATGCTCCACGCCCAGTCGCGTACCGTCTTCATGGGAGAAAATAATACTGTTGATGGGTGTCTGGTCAGAGACATCAAGAAATAACGCCGGAACATTAGTGCAGGCAGCTTCCACAGCAATGGCATCCTGGTCATCCAGCGGATAGTTAATGATCAGCCCACTGACGCGTTGCGCGAGAAGATTGTGCACCGCCGCTTTACAGGCTTCGACGCCGCTTCGTTCTACCATCGACACCACCACGCTGGCACCCAGTTGATCGGCGCGAGATTTAATCGCCGCGACAATTTGCGACGGCGCGTGCAGGGCCAGACTGGAGGTGGCAACGCCAATCAGCAACGACTGTTTGCCCGCCAGTTGTTGTGCCACGCGGTTGGGAATGTAATTCAGCTCCGCCATCGCCGCTTCCACTTTTTCCCGCGTTTTCGCAGAAACGTGGCTGGCCTGGTTCACCACGCGGGAAACGGTCTGATAAGAGACACCGGCATACTCTGCGACATCGTATAACGTTACTGGTTTCACATTCACCACCCTGAATTGACTCTCTTCCGGGCGCTATCATGCCATACCGCGAAAGGTTTTGCGCCATTCGATGGTGTCCGGGATCTCGACGCTCTCCCTTATGCGACTCCTGCATTAGGAAGCAGCCCAGTAGTAGGTTGAGGCCGTTGAGCACCGCCGCCGCAAGGAATGGTGCATGCAAGGAGATGGCGCCCAACAGTCCCCCGGCCACGGGGCCTGCCACCATACCCACGCCGAAACAAGCGCTCATGAGCCCGAAGTGGCGAGCCCGATCTTCCCCATCGGTGATGTCGGCGATATAGGCGCCAGCAACCGCACCTGTGGCGCCGGTGATGCCGGCCACGATGCGTCCGGCGTAGAGGATCGAGATCTCGATCCCGCGAAATTAATACGACTCACTATAGGGGAATTGTGAGCGGATAACAATTCCCCTCTAGAAATAATTTTGTTTAACTTTAAGAAGGAGATATACCATGGGCAGCAGCCATCATCATCATCATCACAGCAGCGGCCTGGTGCCGCGCGGCAGCCATATGGCTAGCATGACTGGTGGACAGCAAATGGGTCGCGGATCCGAATTCGAGCTCCGTCGACAAGCTTGCGGCCGCTGTTAAGCGGCCGCACTCGAGCACCACCACCACCACCACTGAGATCCGGCTGCTAACAAAGCCCGAAAGGAAGCTGAGTTGGCTGCTGCCACCGCTGAGCAATAACTAGCATAACCCCTTGGGGCCTCTAAACGGGTCTTGAGGGGTTTTTTGCTGAAAGGAGGAACTATATCCGGATTGGCGAATGGGACGCGCCCTGTAGCGGCGCATTAAGCGCGGCGGGTGTGGTGGTTACGCGCAGCGTGACCGCTACACTTGCCAGCGCCCTAGCGCCCGCTCCTTTCGCTTTCTTCCCTTCCTTTCTCGCCACGTTCGCCGGCTTTCCCCGTCAAGCTCTAAATCGGGGGCTCCCTTTAGGGTTCCGATTTAGTGCTTTACGGCACCTCGACCCCAAAAAACTTGATTAGGGTGATGGTTCACGTAGTGGGCCATCGCCCTGATAGACGGTTTTTCGCCCTTTGACGTTGGAGTCCACGTTCTTTAATAGTGGACTCTTGTTCCAAACTGGAACAACACTCAACCCTATCTCGGTCTATTCTTTTGATTTATAAGGGATTTTGCCGATTTCGGCCTATTGGTTAAAAAATGAGCTGATTTAACAAAAATTTAACGCGAATTTTAACAAAATATTAACGCTTACAATTTAGGTGGCACTTTTCGGGGAAATGTGCGCGGAACCCCTATTTGTTTATTTTTCTAAATACATTCAAATATGTATCCGCTCATGAATTAATTCTTAGAAAAACTCATCGAGCATCAAATGAAACTGCAATTTATTCATATCAGGATTATCAATACCATATTTTTGAAAAAGCCGTTTCTGTAATGAAGGAGAAAACTCACCGAGGCAGTTCCATAGGATGGCAAGATCCTGGTATCGGTCTGCGATTCCGACTCGTCCAACATCAATACAACCTATTAATTTCCCCTCGTCAAAAATAAGGTTATCAAGTGAGAAATCACCATGAGTGACGACTGAATCCGGTGAGAATGGCAAAAGTTTATGCATTTCTTTCCAGACTTGTTCAACAGGCCAGCCATTACGCTCGTCATCAAAATCACTCGCATCAACCAAACCGTTATTCATTCGTGATTGCGCCTGAGCGAGACGAAATACGCGATCGCTGTTAAAAGGACAATTACAAACAGGAATCGAATGCAACCGGCGCAGGAACACTGCCAGCGCATCAACAATATTTTCACCTGAATCAGGATATTCTTCTAATACCTGGAATGCTGTTTTCCCGGGGATCGCAGTGGTGAGTAACCATGCATCATCAGGAGTACGGATAAAATGCTTGATGGTCGGAAGAGGCATAAATTCCGTCAGCCAGTTTAGTCTGACCATCTCATCTGTAACATCATTGGCAACGCTACCTTTGCCATGTTTCAGAAACAACTCTGGCGCATCGGGCTTCCCATACAATCGATAGATTGTCGCACCTGATTGCCCGACATTATCGCGAGCCCATTTATACCCATATAAATCAGCATCCATGTTGGAATTTAATCGCGGCCTAGAGCAAGACGTTTCCCGTTGAATATGGCTCATAACACCCCTTGTATTACTGTTTATGTAAGCAGACAGTTTTATTGTTCATGACCAAAATCCCTTAACGTGAGTTTTCGTTCCACTGAGCGTCAGACCCCGTAGAAAAGATCAAAGGATCTTCTTGAGATCCTTTTTTTCTGCGCGTAATCTGCTGCTTGCAAACAAAAAAACCACCGCTACCAGCGGTGGTTTGTTTGCCGGATCAAGAGCCACCAACTCTTTTTCCGAAGGTAACTGGCTTCAGCAGAGCGCAGATACCAAATACTGTCCTTCTAGTGTAGCCGTAGTTAGGCCACCACTTCAAGAACTCTGTAGCACCGCCTACATACCTCGCTCTGCTAATCCTGTTACCAGTGGCTGCTGCCAGTGGCGATAAGTCGTGTCTTACCGGGTTGGACTCAAGACGATAGTTACCGGATAAGGCGCAGCGGTCGGGCTGAACGGGGGGTTCGTGCACACAGCCCAGCTTGGAGCGAACGACCTACACCGAACTGAGATACCTACAGCGTGAGCTATGAGAAAGCGCCACGCTTCCCGAAGGGAGAAAGGCGGACAGGTATCCGGTAAGCGGCAGGGTCGGAACAGGAGAGCGCACGAGGGAGCTTCCAGGGGGAAACGCCTGGTATCTTTATAGTCCTGTCGGGTTTCGCCACCTCTGACTTGAGCGTCGATTTTTGTGATGCTCGTCAGGGGGGCGGAGCCTATGGAAAAACGCCAGCAACGCGGCCTTTTTACGGTTCCTGGCCTTTTGCTGGCCTTTTGCTCACATGTTCTTTCCTGCGTTATCCCCTGATTCTGTGGATAACCGTATTACCGCCTTTGAGTGAGCTGATACCGCTCGCCGCAGCCGAACGACCGAGCGCAGCGAGTCAGTGAGCGAGGAAGCGGAAG

2. **Target plasmids and Fragments used in this study**
a. pUC19- γPNA1-151 (γPNA binding sites are highlighted) (3960 bp)

TCGCGCGTTTCGGTGATGACGGTGAAAACCTCTGACACATGCAGCTCCCGGAGACGGTCACAGCTTGTCTGTAAGCGGATGCCGGGAGCAGACAAGCCCGTCAGGGCGCGTCAGCGGGTGTTGGCGGGTGTCGGGGCTGGCTTAACTATGCGGCATCAGAGCAGATTGTACTGAGAGTGCACCAtacAcaGGGTGGTCACGAGGGTGGGCtgActgtacaacaTCCGCATTGAGAACCTCCCTtgAgTATGCGGTGTGAAATACCGCACAGATGCGTAAGGAGAAAATACCGCATCAGGCGCCATTCGCCATTCAGGCTGCGCAACTGTTGGGAAGGGCGATCGGTGCGGGCCTCTTCGCTATTACGCCAGCTGGCGAAAGGGGGATGTGCTGCAAGGCGATTAAGTTGGGTAACGCCAGGGTTTTCCCAGTCACGACGTTGTAAAACGACGGCCAGTGaattcgaacgcgcaatcaaaactaaatacggtagcgataccgagatcaagctcaaatccaaatctgggattatgcatgactccaaatatttggaatcatgggagcggggcagtgcggatatccgtttcgcagagttcgccggcgagaatcgagctcacaacaagcagtttccggctgcgactgtgaatatgggaaggcagccagatggccagggagggatgactcgcgatcgccatgtaagcgttgactacctattgcaaaacctacccaactccccttggacgcaagccttgaaagagggaaagttgtgggatcgagttcaggtccttgctcgcgacggaaaccgttacatgtcaccttcaagactggaatattccgaccccgaacactttacccaactgatggatcaagttggtctgcccgtgtcgatgggtcggcaaagtcatgcgaatagtgtcaagtttgagcagtttgacagacaggcagcggttattgttgcggatggcccgaacttacgtgaggttccagatttgtccccggaaaagttgcaacaactgtctcaaaaagatgtcctgatagcggatcgcaatgaaaaggggcaaagaaccggcacttacactaatgttgtggaatatgagcgcctgatgatgaaattaccgagcgacgcagcgcagcttctcgctgaaccgtccgatagatattcacgtgcttttgtccggccggagccagcattgccccccatcagtgacagccggcggacttatgaaagccgaccgcgcggcccaaccgtaaacagtctgaaaaggccggcggccacgaaaaaggccggccaggcaaaaaagaaaaagtagGCGGCCgcactcgaggcccgaaaggaagctgagttggctgctgccaccgctgagcaataactagcataaccccttggggcctctaaacgggtcttgaggggttttttgctgaaaggaggaactatatccggatatcccgcaagaggcccggcagtaccggcataaccaagcctatgcctacagcatccagggtgacggtgccgaggatgacgatgagcgcattgttagatttcatacacggtgcctgactgcgttagcaatttaactgtgataaactaccgcattaaagcttatcgatgataagctgtcaaacatgagaattcAccTGAGTCCGAGCAGAAGAAGAggAGAGCTCAccGGCTCCCATCACATCAACCggAgGATCCTCTAGAGTCGACCTGCAGGCATGCAAGCTTGGCGTAATCATGGTCATAGCTGTTTCCTGTGTGAAATTGTTATCCGCTCACAATTCCACACAACATACGAGCCGGAAGCATAAAGTGTAAAGCCTGGGGTGCCTAATGAGTGAGCTAACTCACATTAATTGCGTTGCGCTCACTGCCCGCTTTCCAGTCGGGAAACCTGTCGTGCCAGCTGCATTAATGAATCGGCCAACGCGCGGGGAGAGGCGGTTTGCGTATTGGGCGCTCTTCCGCTTCCTCGCTCACTGACTCGCTGCGCTCGGTCGTTCGGCTGCGGCGAGCGGTATCAGCTCACTCAAAGGCGGTAATACGGTTATCCACAGAATCAGGGGATAACGCAGGAAAGAACATGTGAGCAAAAGGCCAGCAAAAGGCCAGGAACCGTAAAAAGGCCGCGTTGCTGGCGTTTTTCCATAGGCTCCGCCCCCCTGACGAGCATCACAAAAATCGACGCTCAAGTCAGAGGTGGCGAAACCCGACAGGACTATAAAGATACCAGGCGTTTCCCCCTGGAAGCTCCCTCGTGCGCTCTCCTGTTCCGACCCTGCCGCTTACCGGATACCTGTCCGCCTTTCTCCCTTCGGGAAGCGTGGCGCTTTCTCATAGCTCACGCTGTAGGTATCTCAGTTCGGTGTAGGTCGTTCGCTCCAAGCTGGGCTGTGTGCACGAACCCCCCGTTCAGCCCGACCGCTGCGCCTTATCCGGTAACTATCGTCTTGAGTCCAACCCGGTAAGACACGACTTATCGCCACTGGCAGCAGCCACTGGTAACAGGATTAGCAGAGCGAGGTATGTAGGCGGTGCTACAGAGTTCTTGAAGTGGTGGCCTAACTACGGCTACACTAGAAGAACAGTATTTGGTATCTGCGCTCTGCTGAAGCCAGTTACCTTCGGAAAAAGAGTTGGTAGCTCTTGATCCGGCAAACAAACCACCGCTGGTAGCGGTGGTTTTTTTGTTTGCAAGCAGCAGATTACGCGCAGAAAAAAAGGATCTCAAGAAGATCCTTTGATCTTTTCTACGGGGTCTGACGCTCAGTGGAACGAAAACTCACGTTAAGGGATTTTGGTCATGAGATTATCAAAAAGGATCTTCACCTAGATCCTTTTAAATTAAAAATGAAGTTTTAAATCAATCTAAAGTATATATGAGTAAACTTGGTCTGACAGTTACCAATGCTTAATCAGTGAGGCACCTATCTCAGCGATCTGTCTATTTCGTTCATCCATAGTTGCCTGACTCCCCGTCGTGTAGATAACTACGATACGGGAGGGCTTACCATCTGGCCCCAGTGCTGCAATGATACCGCGAGACCCACGCTCACCGGCTCCAGATTTATCAGCAATAAACCAGCCAGCCGGAAGGGCCGAGCGCAGAAGTGGTCCTGCAACTTTATCCGCCTCCATCCAGTCTATTAATTGTTGCCGGGAAGCTAGAGTAAGTAGTTCGCCAGTTAATAGTTTGCGCAACGTTGTTGCCATTGCTACAGGCATCGTGGTGTCACGCTCGTCGTTTGGTATGGCTTCATTCAGCTCCGGTTCCCAACGATCAAGGCGAGTTACATGATCCCCCATGTTGTGCAAAAAAGCGGTTAGCTCCTTCGGTCCTCCGATCGTTGTCAGAAGTAAGTTGGCCGCAGTGTTATCACTCATGGTTATGGCAGCACTGCATAATTCTCTTACTGTCATGCCATCCGTAAGATGCTTTTCTGTGACTGGTGAGTACTCAACCAAGTCATTCTGAGAATAGTGTATGCGGCGACCGAGTTGCTCTTGCCCGGCGTCAATACGGGATAATACCGCGCCACATAGCAGAACTTTAAAAGTGCTCATCATTGGAAAACGTTCTTCGGGGCGAAAACTCTCAAGGATCTTACCGCTGTTGAGATCCAGTTCGATGTAACCCACTCGTGCACCCAACTGATCTTCAGCATCTTTTACTTTCACCAGCGTTTCTGGGTGAGCAAAAACAGGAAGGCAAAATGCCGCAAAAAAGGGAATAAGGGCGACACGGAAATGTTGAATACTCATACTCTTCCTTTTTCAATATTATTGAAGCATTTATCAGGGTTATTGTCTCATGAGCGGATACATATTTGAATGTATTTAGAAAAATAAACAAATAGGGGTTCCGCGCACATTTCCCCGAAAAGTGCCACCTGACGTCTAAGAAACCATTATTATCATGACATTAACCTATAAAAATAGGCGTATCACGAGGCCCTTTCGTC

b. pUC19-γPNA1&1-245 (γPNA binding sites are highlighted) (3798 bp)

TCGCGCGTTTCGGTGATGACGGTGAAAACCTCTGACACATGCAGCTCCCGGAGACGGTCACAGCTTGTCTGTAAGCGGATGCCGGGAGCAGACAAGCCCGTCAGGGCGCGTCAGCGGGTGTTGGCGGGTGTCGGGGCTGGCTTAACTATGCGGCATCAGAGCAGATTGTACTGAGAGTGCACCATACACAGGGTGGTCACGAGGGTGGGCTGACTGTACAACATCCGCATTGAGAACCTCCCTTGAGTATGCGGTGTGAAATACCGCACAGATGCGTAAGGAGAAAATACCGCATCAGGCGCCATTCGCCATTCAGGCTGCGCAACTGTTGGGAAGGGCGATCGGTGCGGGCCTCTTCGCTATTACGCCAGCTGGCGAAAGGGGGATGTGCTGCAAGGCGATTAAGTTGGGTAACGCCAGGGTTTTCCCAGTCACGACGTTGTAAAACGACGGCCAGTGAATTCGAACGCGCAATCAAAACTAAATACGGTAGCGATACCGAGATCAAGCTCAAATCCAAATCTGGGATTATGCATGACTCCAAATATTTGGAATCATGGGAGCGGGGCAGTGCGGATATCCGTTTCGCAGAGTTCGCCGGCGAGAATCGAGCTCACAACAAGCAGTTTCCGGCTGCGACTGTGAATATGGGAAGGCAGCCAGATGGCCAGGGAGGGATGACTCGCGATCGCCATGTAAGCGTTGACTACCTATTGCAAAACCTACCCAACTCCCCTTGGACGCAAGCCTTGAAAGAGGGAAAGTTGTGGGATCGAGTTCAGGTCCTTGCTCGCGACGGAAACCGTTACATGTCACCTTCAAGACTGGAATATTCCGACCCCGAACACTTTACCCAACTGATGGATCAAGTTGGTCTGCCCGTGTCGATGGGTCGGCAAAGTCATGCGAATAGTGTCAAGTTTGAGCAGTTTGACAGACAGGCAGCGGTTATTGTTGCGGATGGCCCGAACTTACGTGAGGTTCCAGATTTGTCCCCGGAAAAGTTGCAACAACTGTCTCAAAAAGATGTCCTGATAGCGGATCGCAATGAAAAGGGGCAAAGAACCGGCACTTACACTAATGTTGTGGAATATGAGCGCCTGATGATGAAATTACCGAGCGACGCAGCGCAGCTTCTCGCTGAACCGTCCGATAGATATTCACGTGCTTTTGTCCGGCCGGAGCCAGCATTGCCCCCCATCAGTGACAGCCGGCGGACTTATGAAAGCCGACCGCGCGGCCCAACCGTAAACAGTCTGAAAAGGCCGGCGGCCACGAAAAAGGCCGGCCAGGCAAAAAAGAAAAAGTAGGCGGCCGCACTCGAGCAATTGTCCCCCCCTTCCCTCCCACCCCCTGCCAAGTCTCCCTCCCAGGATCTTCTCTGGCTCCATCGTAAGCAAACCTTAGAGGTTCTGGCAAGGAGAGAGATGGGAGCTCCCATGGACACAGTTGAAGGAAGGAAAGATGATTCCTGGGAGAGATCTTCTTCTGCTCGGACTCAGGGTGGTCACGAGGGTGGGCGCATGCAGGATCCTCTAGAGTCGACCTGCAGGCATGCAAGCTTGGCGTAATCATGGTCATAGCTGTTTCCTGTGTGAAATTGTTATCCGCTCACAATTCCACACAACATACGAGCCGGAAGCATAAAGTGTAAAGCCTGGGGTGCCTAATGAGTGAGCTAACTCACATTAATTGCGTTGCGCTCACTGCCCGCTTTCCAGTCGGGAAACCTGTCGTGCCAGCTGCATTAATGAATCGGCCAACGCGCGGGGAGAGGCGGTTTGCGTATTGGGCGCTCTTCCGCTTCCTCGCTCACTGACTCGCTGCGCTCGGTCGTTCGGCTGCGGCGAGCGGTATCAGCTCACTCAAAGGCGGTAATACGGTTATCCACAGAATCAGGGGATAACGCAGGAAAGAACATGTGAGCAAAAGGCCAGCAAAAGGCCAGGAACCGTAAAAAGGCCGCGTTGCTGGCGTTTTTCCATAGGCTCCGCCCCCCTGACGAGCATCACAAAAATCGACGCTCAAGTCAGAGGTGGCGAAACCCGACAGGACTATAAAGATACCAGGCGTTTCCCCCTGGAAGCTCCCTCGTGCGCTCTCCTGTTCCGACCCTGCCGCTTACCGGATACCTGTCCGCCTTTCTCCCTTCGGGAAGCGTGGCGCTTTCTCATAGCTCACGCTGTAGGTATCTCAGTTCGGTGTAGGTCGTTCGCTCCAAGCTGGGCTGTGTGCACGAACCCCCCGTTCAGCCCGACCGCTGCGCCTTATCCGGTAACTATCGTCTTGAGTCCAACCCGGTAAGACACGACTTATCGCCACTGGCAGCAGCCACTGGTAACAGGATTAGCAGAGCGAGGTATGTAGGCGGTGCTACAGAGTTCTTGAAGTGGTGGCCTAACTACGGCTACACTAGAAGAACAGTATTTGGTATCTGCGCTCTGCTGAAGCCAGTTACCTTCGGAAAAAGAGTTGGTAGCTCTTGATCCGGCAAACAAACCACCGCTGGTAGCGGTGGTTTTTTTGTTTGCAAGCAGCAGATTACGCGCAGAAAAAAAGGATCTCAAGAAGATCCTTTGATCTTTTCTACGGGGTCTGACGCTCAGTGGAACGAAAACTCACGTTAAGGGATTTTGGTCATGAGATTATCAAAAAGGATCTTCACCTAGATCCTTTTAAATTAAAAATGAAGTTTTAAATCAATCTAAAGTATATATGAGTAAACTTGGTCTGACAGTTACCAATGCTTAATCAGTGAGGCACCTATCTCAGCGATCTGTCTATTTCGTTCATCCATAGTTGCCTGACTCCCCGTCGTGTAGATAACTACGATACGGGAGGGCTTACCATCTGGCCCCAGTGCTGCAATGATACCGCGAGACCCACGCTCACCGGCTCCAGATTTATCAGCAATAAACCAGCCAGCCGGAAGGGCCGAGCGCAGAAGTGGTCCTGCAACTTTATCCGCCTCCATCCAGTCTATTAATTGTTGCCGGGAAGCTAGAGTAAGTAGTTCGCCAGTTAATAGTTTGCGCAACGTTGTTGCCATTGCTACAGGCATCGTGGTGTCACGCTCGTCGTTTGGTATGGCTTCATTCAGCTCCGGTTCCCAACGATCAAGGCGAGTTACATGATCCCCCATGTTGTGCAAAAAAGCGGTTAGCTCCTTCGGTCCTCCGATCGTTGTCAGAAGTAAGTTGGCCGCAGTGTTATCACTCATGGTTATGGCAGCACTGCATAATTCTCTTACTGTCATGCCATCCGTAAGATGCTTTTCTGTGACTGGTGAGTACTCAACCAAGTCATTCTGAGAATAGTGTATGCGGCGACCGAGTTGCTCTTGCCCGGCGTCAATACGGGATAATACCGCGCCACATAGCAGAACTTTAAAAGTGCTCATCATTGGAAAACGTTCTTCGGGGCGAAAACTCTCAAGGATCTTACCGCTGTTGAGATCCAGTTCGATGTAACCCACTCGTGCACCCAACTGATCTTCAGCATCTTTTACTTTCACCAGCGTTTCTGGGTGAGCAAAAACAGGAAGGCAAAATGCCGCAAAAAAGGGAATAAGGGCGACACGGAAATGTTGAATACTCATACTCTTCCTTTTTCAATATTATTGAAGCATTTATCAGGGTTATTGTCTCATGAGCGGATACATATTTGAATGTATTTAGAAAAATAAACAAATAGGGGTTCCGCGCACATTTCCCCGAAAAGTGCCACCTGACGTCTAAGAAACCATTATTATCATGACATTAACCTATAAAAATAGGCGTATCACGAGGCCCTTTCGTC

**Supplementary tables**

**Table S1:** Different proteins sequences employed in this study

| **Protein name** | **Sequence** |
| --- | --- |
| (wt)RusA | MNTYSITLPWPPSNNRYYRHNRGRTHVSAEGQAYRDNVARIIKNAMLDIGLAMPVKIRIECHMPDRRRRDLDNLQKAAFDALTKAGFWLDDAQVVDYRVVKMPVTKGGRLELTITEMGNE |
| (d)RusA | MNTYSITLPWPPSNNRYYRHNRGRTHVSAEGQAYRDNVARIIKNAMLDIGLAMPVKIRIECHMPDRRRRNLDNLQKAAFDALTKAGFWLDDAQVVDYRVVKMPVTKGGRLELTITEMGNE |
| FokI-(d)RusA | MQLVKSELEEKKSELRHKLKYVPHEYIELIEIARNSTQDRILEMKVMEFFMKVYGYRGKHLGGSRKPDGAIYTVGSPIDYGVIVDTKAYSGGYNLPIGQADEMQRYVEENQTRNKHINPNEWWKVYPSSVTEFKFLFVSGHFKGNYKAQLTRLNHITNCNGAVLSVEELLIGGEMIKAGTLTLEEVRRKFNNGEINFSGGSSGGSSGSETPGTSESATPESSGGSSGGSSMNTYSITLPWPPSNNRYYRHNRGRTHVSAEGQAYRDNVARIIKNAMLDIGLAMPVKIRIECHMPDRRRRNLDNLQKAAFDALTKAGFWLDDAQVVDYRVVKMPVTKGGRLELTITEMGNE |
| (d)RusA-FokI | MNTYSITLPWPPSNNRYYRHNRGRTHVSAEGQAYRDNVARIIKNAMLDIGLAMPVKIRIECHMPDRRRRNLDNLQKAAFDALTKAGFWLDDAQVVDYRVVKMPVTKGGRLELTITEMGNESGGSSGGSSGSETPGTSESATPESSGGSSGGSSQLVKSELEEKKSELRHKLKYVPHEYIELIEIARNSTQDRILEMKVMEFFMKVYGYRGKHLGGSRKPDGAIYTVGSPIDYGVIVDTKAYSGGYNLPIGQADEMQRYVEENQTRNKHINPNEWWKVYPSSVTEFKFLFVSGHFKGNYKAQLTRLNHITNCNGAVLSVEELLIGGEMIKAGTLTLEEVRRKFNNGEINF |

**Table S2:** Lysis buffer composition

| **Component** | **Stock** | **Concentration (mM) / Amount per 50 mL** |
| --- | --- | --- |
| Tris–HCl pH7.5 | 1 M | 50 mM |
| NaCl | 5 M | 300 mM |
| Glycerol | 50% | 5% |
| Thermo Scientific, 20491 | Powder | 1 mM |
| PMSF | 100 mM | 0.5 mM |
| Lysozyme (Sigma, L6876) | Powder | 100 mg |
| MgCl2 | 1 M | 4.5 mM |
| Protease inhibitor tablet (Thermo Scientific, A32953) | Tablets | 1 tablet |
| Imidazole | 3 M | 20 mM |
| Benzonase (Merck, E1014-5KU) | 25 units/µL | 12.5 units |
| NP-40 | 10% | 0.1% |
| 2-Mercaptoethanol (Aldrich, M6250) | 14.3 M | 4.86 mM |

**Table S3:** Sequences of oligos used to prepare Holliday Junctions

| **Strand** | **Strand Sequence 5’ to 3’** |
| --- | --- |
| S1 | AGCTGATCGTACGATGCTAGTCGTCCATCAGGCTAGTCGATGCTAGCTGACGTAGCTGATCGTAGCATCGTGCAGCTGATCGTAGCATCCGAATCA |
| S2 | ACGATGCTCGATCGATGCATGCTAGTCGACGTCGATCGTACGATGCTAGTCGATGCTAGCTGAGCTGCTAGGACGACTAGCATCGTACGATCAGCT |
| S3 | GCTAGCTAGTCGATCGTAGCATATCCTAGCAGCTCAGCTAGCATCGACTAGCATCGTACGATCGACGTCGACTAGCATGCATCGATCGAGCATCGT |
| S4 | TGATTCGGATGCTACGATCAGCTGCACGATGCTACGATCAGCTACGTCAGCTAGCATCGACTAGCCTGATGGATATGCTACGATCGACTAGCTAGC |

Homology core sequence on each strand is highlighted in red

Digestion sites for PuvII and SphI are highlighted in yellow and green, respectively

**Table S4:** γPNA1 sequence tested in the current study

| **PNA** | **Sequence** | **PNA modifications** | **Tm Values** |
| --- | --- | --- | --- |
| γPNA1 | H-KKK-GCCCACCCTCGTGACCACCC- KKK-propargylglycine-NH_2_ | Gamma-alanine at all bases | 85.0°C |

H, free amine at the N terminus.

NH2, amide at the C terminus.

K, lysine.

**Table S5:** Top and bottom oligos used to clone the target sequences in pUC19 vectors

| **Oligo name** | **Oligo sequence (5’---3’)** |
| --- | --- |
| 151_ γPNA1_top with NdeI | TACACAGGGTGGTCACGAGGGTGGGCTGACTGTACAACATCCGCATTGAGAACCTCCCTTGAG |
| 151_ γPNA1_bottom with NdeI | TACTCAAGGGAGGTTCTCAATGCGGATGTTGTACAGTCAGCCCACCCTCGTGACCACCCTGTG |

**Table S6:** Primers used in the current study

| **Primer name** | **Prime Sequence (5’----3’)** | **Employed for** |
| --- | --- | --- |
| pET28 seq F | GCATGACTGGTGGACAGCAA | Sanger sequencing of pET28A-RusA expression clones |
| pET28 seq R | AGAGGCCCCAAGGGGTTATG |  |
| 151-pUC19_PNA1_F | TGTCTGTAAGCGGATGCCGG | All the pUC19 target plasmids confirmation by Sanger sequencing, PCR amplification of pUC19 targets for producing the |
| 151-pUC19_PNA1_R | GCCAGCTGGCGTAATAGCG |  |

**Table S7:** Compositions of buffers used in this study

| **Buffer** | **Composition 1X** |
| --- | --- |
| **NEB rCutSmart** \| | 50 mM Potassium Acetate 20 mM Tris-acetate 10 mM Magnesium Acetate 100 μg/ml Recombinant Albumin pH 7.9@25°C |
| **NEB r1.1** \| | 10 mM Bis-Tris-Propane-HCl 10 mM MgCl2 100 μg/ml Recombinant Albumin pH 7.0@25°C |
| **NEB r2.1** \| | 50 mM NaCl 10 mM Tris-HCl 10 mM MgCl2 100 μg/ml Recombinant Albumin pH 7.9@25°C |
| **NEB r3.1** \| | 100 mM NaCl 50 mM Tris-HCl 10 mM MgCl2 100 μg/ml Recombinant Albumin pH 7.9@25°C |
| **HEPES Buffer** \| | 1M HEPES  pH 7.5 |
| **MOPS Buffer** \| | 20 mM MOPS  5 mM CH3COONa  1 mM EDTA  pH 7.0 |
| **PBS Buffer** | 137 mM NaCl 2.7 mM KCl 10 mM Na₂HPO₄ 1.8 mM KH₂PO₄ pH 7.4 |

NEB Buffers composition adapted from (<https://www.bioke.com/webshop/neb/b7030.html>)
